# Stable antibiotic resistance and rapid human adaptation in livestock-associated MRSA

**DOI:** 10.1101/2021.08.20.457141

**Authors:** Marta Matuszewska, Gemma G. R. Murray, Xiaoliang Ba, Rhiannon Wood, Mark A. Holmes, Lucy A. Weinert

## Abstract

Mobile genetic elements (MGEs) are agents of horizontal gene transfer in bacteria, but can also be vertically inherited by daughter cells. Establishing the dynamics that led to contemporary patterns of MGEs in bacterial genomes is central to predicting the emergence and evolution of novel and resistant pathogens. Methicillin-resistant *Staphylococcus aureus* (MRSA) clonal-complex (CC) 398 is the dominant MRSA in European livestock and a growing cause of human infections. Previous studies have identified three categories of MGEs whose presence or absence distinguishes livestock-associated CC398 from a closely related and less antibiotic-resistant human-associated population. Here we fully characterise the evolutionary dynamics of these MGEs using a collection of 1,180 CC398 genomes, sampled from livestock and humans, over 27 years. We find that the emergence of livestock-associated CC398 coincided with the acquisition of a Tn*916* transposon carrying a tetracycline resistance gene, which has been stably inherited for 57 years. This was followed by the acquisition of a type V SCC*mec* that carries methicillin, tetracycline and heavy metal resistance genes, which has been maintained for 35 years, with occasional truncations and replacements with type IV SCC*mec*. In contrast, a class of prophages that carry a human immune evasion gene cluster and that are largely absent from livestock-associated CC398, have been repeatedly gained and lost in both human- and livestock-associated CC398. These contrasting dynamics mean that when livestock-associated MRSA is transmitted to humans, adaptation to the human host outpaces loss of antibiotic resistance. In addition, the stable inheritance of resistance-associated MGEs suggests that the impact of ongoing reductions in antibiotic and zinc oxide use in European farms on livestock-associated MRSA will be slow to be realised.

## Introduction

Mobile genetic elements (MGEs) play an important role in the evolution of bacterial pathogens. They can move rapidly between bacterial genomes, but can also be vertically inherited through stable integration into a host genome. As MGEs often carry genes associated with virulence and antibiotic resistance (Frost et al. 2005; Rankin et al. 2011), an understanding of the drivers and barriers to their acquisition and maintenance is central to predicting the emergence and evolution of novel and resistant pathogens (Brockhurst et al. 2019).

The emergence and evolution of methicillin-resistant *Staphylococcus aureus (*MRSA) across different ecological niches and host species is associated with the horizontal transfer of MGEs. Methicillin resistance is carried by the staphylococcal cassette chromosome element SCC*mec*, and additional MGEs carry resistance to other antibiotics, virulence factors and host-specific adaptations (Hanssen and Ericson Sollid 2006; Jamrozy et al. 2017; Haag et al. 2019; Turner et al. 2019; Matuszewska et al. 2020). While most MRSA clonal complexes (CCs) show an association with specific MGEs, their dynamics are not widely understood, leading to a gap in our understanding of the adaptive potential of *S. aureus* CCs.

Intensification combined with high levels of antibiotic use in farming has led to particular concerns about livestock as reservoirs of antibiotic-resistant human infections(WHO 2015). CC398 has become the dominant MRSA in European livestock. Its rise has been particularly evident in Danish pig farms where the proportion of MRSA-positive herds has increased from <5% in 2008 to 90% in 2018 (Sieber et al. 2018; DANMAP 2019), but it has also been observed in other European countries, and other livestock species (Lekkerkerk et al. 2015; Islam et al. 2017; Anjum et al. 2019). Livestock-associated (LA) MRSA CC398 has been associated with increasing numbers of human infections, in both people with and without direct contact with livestock (Larsen et al. 2017; van Alen et al. 2017; Sieber et al. 2019). Understanding the emergence and success of CC398 in European livestock and its capacity to infect the human host is integral to managing the risk that it, and other livestock-associated pathogens, pose to public health.

Previous studies have used genome sequences to reconstruct the evolutionary history of CC398 (Price et al. 2012; Ward et al. 2014; Silva et al. 2017). They identified a largely methicillin-resistant and livestock-associated clade of CC398, that falls as either sister to (Ward et al. 2014) or within (Price et al. 2012; Silva et al. 2017) a largely methicillin-sensitive and human-associated clade. Through comparing the genomes of isolates from livestock-associated and human-associated CC398, these studies concluded that the emergence of CC398 in livestock was associated with both the acquisition of antibiotic resistance genes and the loss of genes associated with human immune evasion. While these genes are known to be carried on three categories of MGEs (Tn*916* conjugative transposons, SCC*mec* and ϕSa3 prophages), little is known about the dynamics of these MGEs within CC398. Here, we undertake a comprehensive reconstruction of the evolutionary dynamics of these MGEs. We find that while their patterns of presence/absence all show a strong association with the transition to livestock, this is the result of contrasting dynamics. These dynamics can inform predictions about the risk posed by LA-MRSA spillover infections in humans, and the resilience of antibiotic resistance in LA-MRSA to ongoing changes in antibiotic use in farming.

## Results

### Livestock-associated CC398 emerged between 1957 and 1970

We collected and assembled publicly available whole-genome sequence data from CC398, and sequenced five isolates recently sampled from pig farms in the UK. Our collection includes high-quality whole genome assemblies of 1,180 isolates (including 43 complete reference genomes). This collection spans 15 host species (including humans, pigs, cows, chickens, turkeys and horses), 28 countries (across Europe, America, Asia, and Australasia), and 27 years (1992 to 2018) (Figure 1-figure supplement 1, Supplementary File 1).

We constructed a recombination-stripped maximum likelihood phylogeny of CC398 using reference-mapped assemblies of our collection. We rooted the phylogeny with outgroups from four other *S. aureus* sequence types (STs) in four separate reconstructions. We also constructed a phylogeny from a recombination-stripped concatenated alignment of core genes extracted from *de novo* assemblies, with a midpoint rooting. These reconstructions consistently returned the same topology, and one that is described in previous studies: a livestock-associated clade of CC398 (704 isolates) that falls within a more diverse, and other largely human-associated clade (476 isolates) (Figure 1a and Figure 1-figure supplement 2) (Price et al. 2012; Silva et al. 2017).

**Figure 1.**
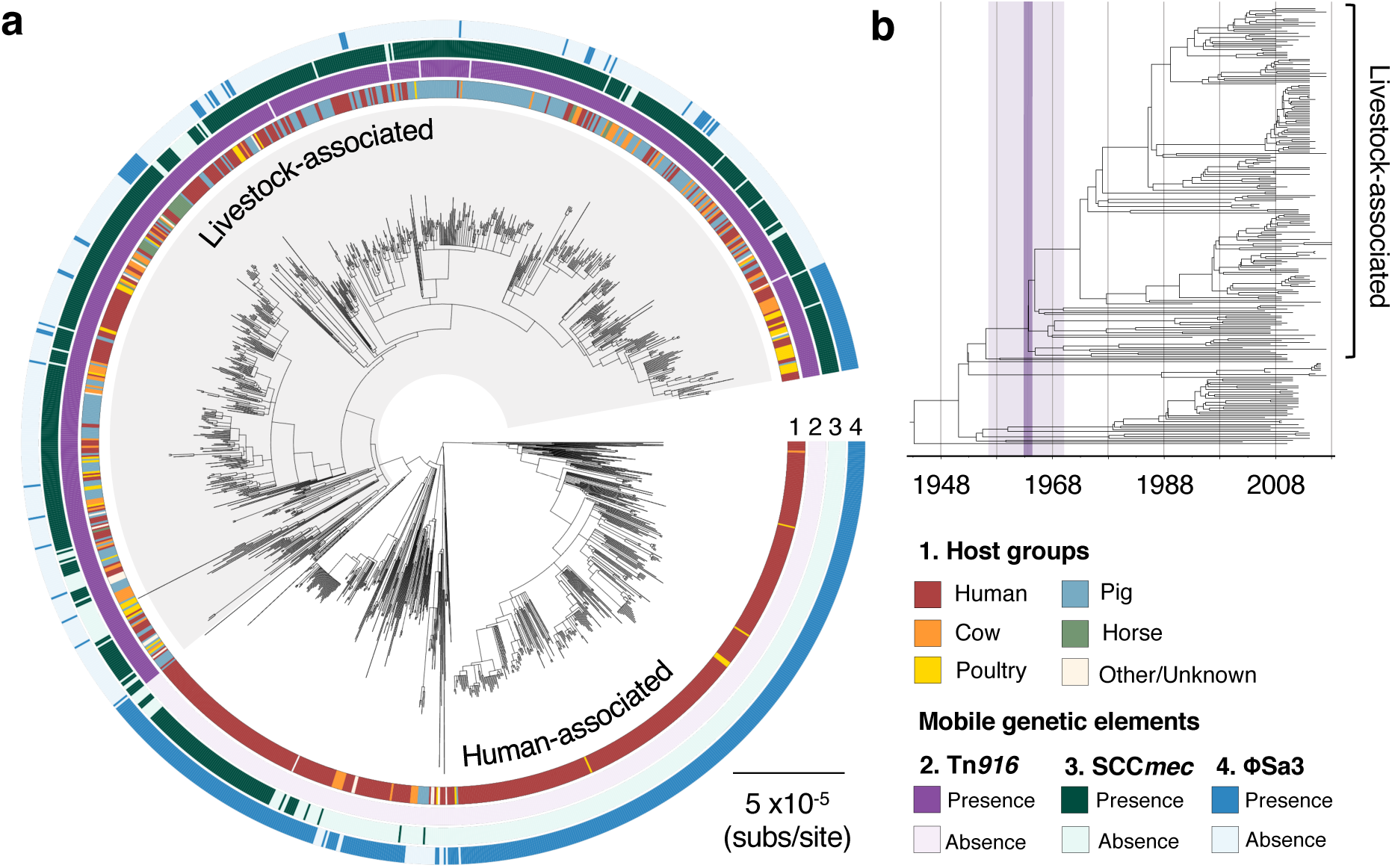
The transition to livestock-association in the 1960s was accompanied by changes in the frequencies of three mobile genetic elements. (a) A maximum likelihood phylogeny of 1,180 isolates of CC398, rooted using an outgroup from ST291. Grey shading indicates the livestock-associated clade. Outer rings describe (1) the host groups isolates were sampled from, and the presence of three MGEs: (2) a Tn*916* transposon carrying *tetM*, (3) a SCC*mec* carrying *mecA*, and (4) a ϕSa3 prophage carrying a human immune evasion gene cluster. (b) A dated phylogeny of a sample of 250 CC398 isolates, that shows livestock-associated CC398 originated around 1964 (95% HPD: 1957-1970).

We used the temporal structure in our collection to date the origin of the livestock-associated clade. Due to the size of our collection, we constructed dated phylogenies from three subsampled data sets, each of which includes 250 isolates. Our estimates of the evolutionary rate were consistent across all reconstructions (1.1-1.6 ×10^−6^ subs/site/year), and similar to estimates from previous studies of CC398 (Ward et al. 2014) and other *S. aureus* CCs (Hsu et al. 2015). This led to an estimate of the origin of the livestock-associated clade of approximately 1964 (95% CI: 1957-1970) (Figure 1b, Figure 1-figure supplement 4 and Figure 1- figure supplement 4).

### The transition to livestock-association is associated with changes in the frequencies of three mobile genetic elements with very different dynamics

Comparisons of the genomes of isolates from human- and livestock-associated CC398, have previously indicated that the transition to livestock was associated with the acquisition of genes associated with both tetracycline and methicillin resistance (*tetM* and *mecA*), and the loss of genes associated with human immune evasion (the immune evasion gene cluster) (Price et al. 2012). Our analyses of this larger collection are broadly consistent with this. We find that the genes whose presence most strongly distinguishes isolates from the human-and livestock-associated groups are associated with three categories of MGEs: (1) a Tn*916* transposon carrying *tetM*; (2) SCC*mec* carrying *mecA*; and (3) ϕSa3 prophages carrying a human immune evasion gene cluster (Figure 1a, Figure 1-figure supplement 5, Supplementary File 2). Genes associated with the Tn*916* transposon and SCC*mec* elements are more common in the livestock-associated clade, while the reverse is true of genes associated with ϕSa3 prophages.

#### 1. Stable maintenance of a Tn916 transposon carrying tetM

We identified a contiguous assembly of a Tn*916* transposon carrying *tetM* in 699/704 isolates in our collection of livestock-associated CC398 (Figure 2a, Supplementary File 3 and Supplementary File 4) (de Vries et al. 2009; Roberts and Mullany 2009). Several lines of evidence indicate that the presence of the Tn*916* transposon in livestock-associated CC398 is the result of a single acquisition event, followed by stable inheritance. First, the location of the transposon in the genome of livestock-associated CC398 is conserved (Tn*916* is always found next to the same core gene; WP_000902814 in the published annotation of S0385). Second, an alignment of the coding regions of this element (extracted from *de novo* assemblies) shows a similar average nucleotide diversity to core genes (Figure 2d). Third, a phylogeny constructed from the genes in this element is entirely congruent with the phylogeny of livestock-associated CC398 (Figure 2b,c).

**Figure 2.**
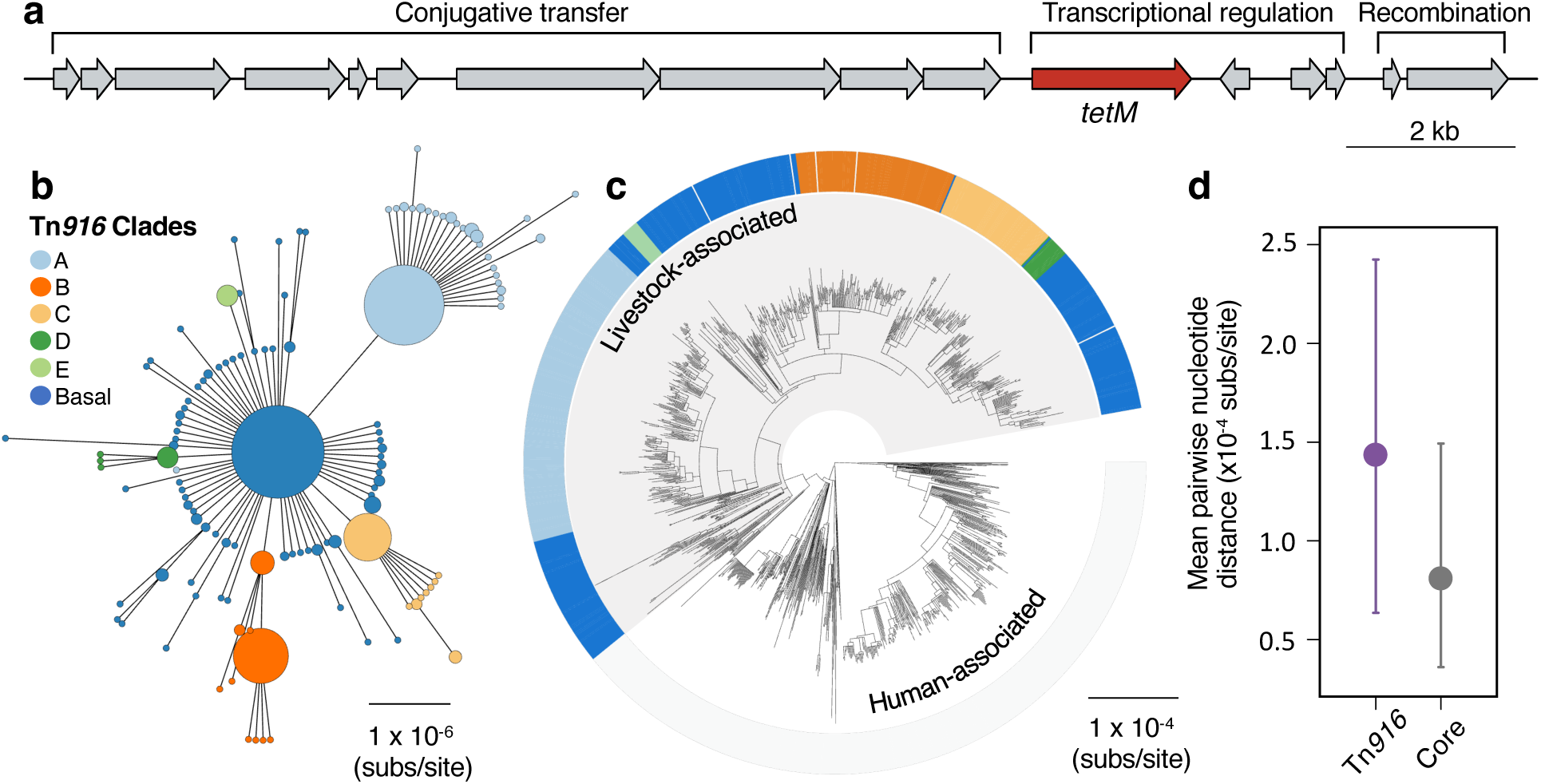
A Tn*916* transposon carrying *tetM* has been stably maintained by livestock-associated CC398 since its origin. (a) A gene map of the Tn*916* transposon in CC398 (based on reference genome 1_1439), with annotations based on previous studies(de Vries et al. 2009; Roberts and Mullany 2009). (b) A minimum spanning tree of the element based on a concatenated alignment of all genes shown in (a). Points represent groups of identical elements, with point size correlated with number of elements on a log-scale, and colours representing well-supported clades (>70 bootstrap support in a maximum likelihood phylogeny) that include >10 elements (smaller clades are incorporated into their basal clade). (c) These clades are annotated onto the CC398 phylogeny as an external ring. (d) Mean pairwise nucleotide distance between isolates carrying the Tn*916* transposon based on genes in the Tn*916* transposon and core genes, using bootstrapping to estimate error (see methods for details).

Our analyses indicate that the Tn*916* transposon has been maintained in livestock-associated CC398 since its origin, and therefore for around 57 years (Figure 1b). Nevertheless, its absence from 5/704 livestock-associated CC398 isolates in our collection suggests that it remains capable of excision (this is consistent with experimental studies that have shown that Tn*916* in CC398 is a functional conjugative transposon; de Vries et al. 2009). None of the genes associated with this element are present in these five isolates, and in two we were able to identify an intact integration site in the assembled genome (Figure 2-figure supplement 1). These five isolates are broadly distributed across the livestock-associated clade, and not linked to a particular host species or geographic location.

#### 2. More variable maintenance of a SCCmec carrying mecA, tetK and czrC

Previous studies suggest that LA-MRSA CC398 emerged from human-associated MSSA (Price et al. 2012). However, the presence of SCC*mec* elements in recently sampled human-associated CC398 isolates that fall basal to the livestock-associated clade, including clinical isolates from China (He et al. 2018; Zou et al. 2022), Denmark (Møller et al. 2019), and New Zealand (Silva et al. 2017), makes the association between methicillin-resistance and livestock-association less clear (Figure 1a).

SCC*mec* elements in *S. aureus* are categorised into several types (Hanssen and Ericson Sollid 2006). Consistent with previous studies of CC398 (e.g. Price et al. 2012), we observe both type V (76%) and type IV (21%) in CC398 (we were unable to confidently type the remaining 3%; Figure 3c, Figure 3-figure supplement 1, Supplementary File 3 and Supplementary File 5). Most of the type V SCC*mec* elements belong to the subtype Vc previously described in livestock-associated CC398 (Li et al. 2011; Price et al. 2012; Vandendriessche et al. 2014). This element includes two additional resistance genes: *tetK* (tetracycline resistance) and *czrC* (heavy metal resistance) (Figure 3a, Supplementary File 6). We identified a full-length version of this element in 335 genomes (including some of the most recent isolates in our collection, sampled from UK pig farms in 2018) and shorter type V elements in 204 genomes. Full-length versions are only observed in the livestock-associated clade, while shorter versions are found in both livestock-and human-associated groups of CC398. Shorter versions often lacked *tetK* (n=90) and *czrC* (n=117) genes (Figure 3-figure supplement 2). Type IV elements are only observed in the livestock-associated clade. They include subtypes IVa and IVc, and all only carry a single resistance gene: *mecA*.

**Figure 3.**
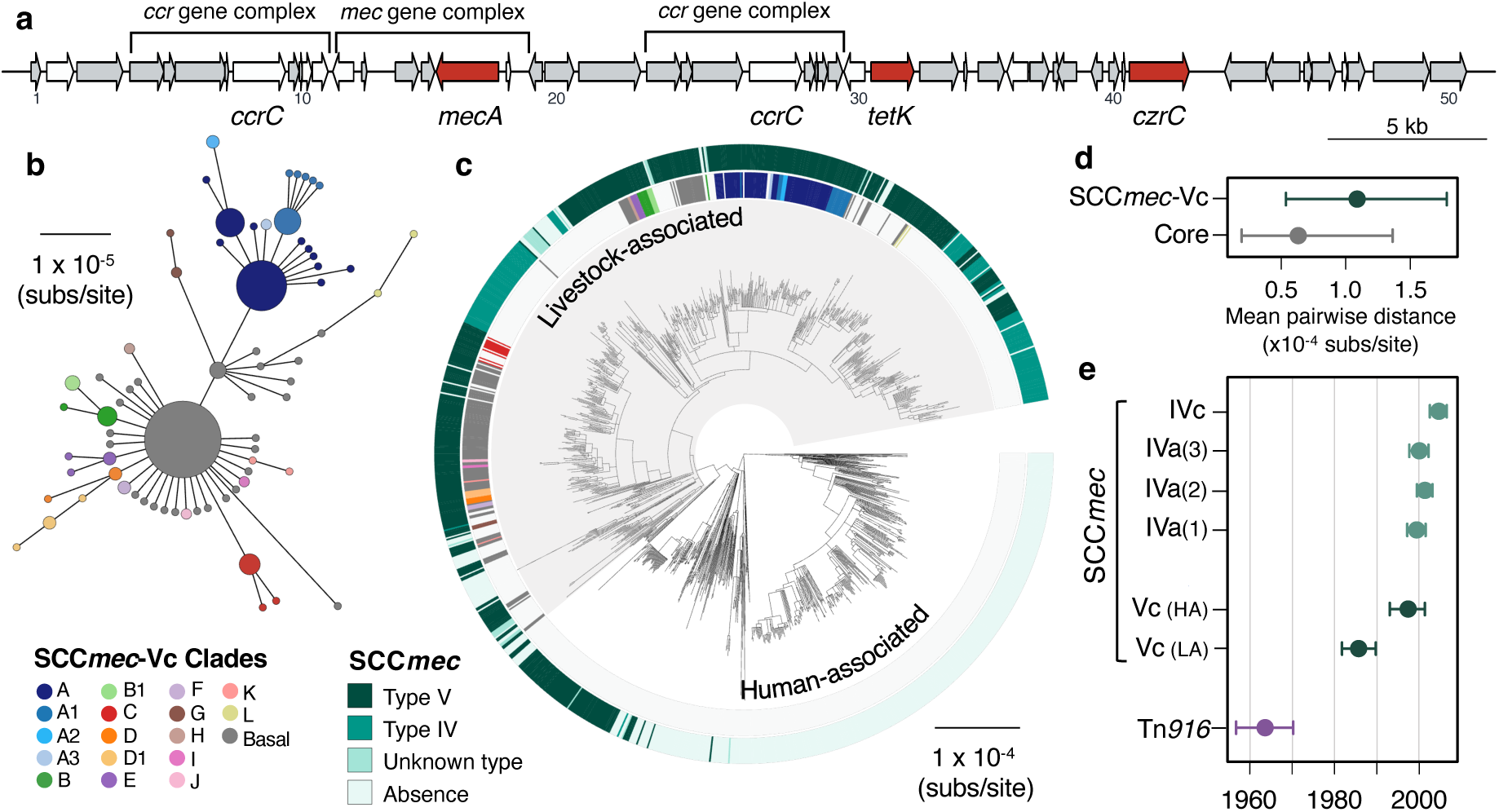
A type V SCC*mec* has been maintained since the 1980s, with occasional replacements. (a) A gene map of the type Vc *SCCmec* element in CC398 (using the 1_1439 reference strain), with annotations from previous studies (Li et al. 2011; Vandendriessche et al. 2014). Genes in white were excluded from analyses of diversity within the element due to difficulties in distinguishing homologues. (b) A minimum-spanning tree of the type Vc *SCCmec* element based on a concatenated alignment of the genes (grey and red) in (a). Points represent groups of identical elements, point size correlates with group size on a log-scale, and colours represent well-supported clades (>70 bootstrap support in a maximum likelihood phylogeny). (c) Well-supported clades and SCC*mec* type are annotated on the CC398 phylogeny in external rings. (d) Mean pairwise nucleotide distance between isolates carrying the SCC*mec* type Vc based on genes in the SCC*mec* type Vc and core genes, with error estimated by bootstrapping (see methods for details). (e) Acquisition dates for different SCC*mec* elements and Tn*916* inferred from an ancestral state reconstruction over the dated phylogeny in 1 (b). Dates for type Vc are shown for both livestock and human-associated CC398.

While type Vc is the most common SCC*mec* in our collection of livestock-associated CC398, this largely reflects isolates from pigs. Type Vc SCC*mec* is much more common in pigs (77% of isolates; n=286) than type IV SCC*mec* (7% of isolates). In cows (n=74) the difference is reduced: 45% are type Vc and 32% are type IV. And in isolates from other animal species (n=94) type IV elements (55%) are more common than type Vc (33%).

Diversity within the type Vc SCC*mec* element indicates that a full-length type Vc SCC*mec* was acquired once by livestock-associated CC398, and has been maintained within CC398 largely through vertical transmission. First, low nucleotide diversity within full-length versions of the element is consistent with 329/335 sharing a recent origin common within livestock-associated CC398 (Figure 3d). Second, patterns of diversity are largely congruent with the core genome phylogeny, consistent with vertical inheritance (Figure 3b,c). Third, low nucleotide diversity within genes shared across full-and shorter-length versions of the element are consistent with most shorter-length versions being the result of deletion within livestock-associated CC398 (Figure 3-figure supplement 2). In contrast, diversity in the type IV elements in livestock-associated CC398 supports four independent acquisitions from outside of CC398 (Figure 3-figure supplement 3). Similar to the type Vc element, once acquired, these elements tend to be maintained.

While the SCC*mec* elements carried by human-associated CC398 are always type V, 70/80 fall within a single clade from a hospital outbreak in Denmark in 2016 (Møller et al. 2019). They show a truncation relative to the full-length type Vc that is also observed in 3 isolates from livestock-associated CC398 (leading to the absence of *czrC*, Figure 3-figure supplement 2). Pairwise nucleotide distances between these 70 human-associated CC398 elements and the full-length type Vc in livestock-associated CC398 are consistent with a recent common ancestor within CC398. In contrast, nucleotide diversity within the other 10 type V SCC*mec* in human-associated CC398 and distances from the livestock-associated CC398 type Vc indicate multiple independent acquisitions from outside of CC398 (Figure 3-figure supplement 2).

Using our dated phylogeny of CC398 and categorisation of SCC*mec* based on diversity within the elements, we inferred the dynamics of gain and loss within CC398 (Figure 3e and Figure 3-figure supplement 4). These reconstructions consistently estimated that the type Vc SCC*mec* had been acquired by livestock-associated CC398 by around 1986 (95% CI: 1982-1990), and has therefore been maintained within livestock-associated CC398 for around 35 years. While the diversity within this element indicates a single acquisition by CC398, these reconstructions indicate multiple gains, likely reflecting horizontal transmission within CC398. They also indicate that the acquisition by livestock-associated CC398 predated the acquisition by human-associated CC398, consistent with transmission from livestock-associated CC398 to human-associated CC398. In contrast, type IV elements show evidence of several more recent acquisitions in relatively quick succession between 1997 and 2004 (Figure 3e and Figure 3-figure supplement 4).

Together, our analyses are consistent with LA-MRSA emerging from human-associated MSSA and acquiring the type Vc SCC*mec* element following its initial diversification. The element has been stably maintained (particularly in pigs), although with several exceptions -including deletions of parts of the element and replacements with smaller type IV SCC*mec*.

#### 3. Loss of a ϕSa3 prophage carrying a human immune evasion gene cluster, but with frequent reacquisition

In contrast to the maintenance of Tn*916* and SCC*mec* type V in the livestock-associated clade, the ϕSa3 prophage is highly dynamic. ϕSa3 prophages carry human immune evasion gene clusters that include a variable set of functional genes that encode human-specific virulence factors, including *sak, chp, scn, sea* and *sep* (Gladysheva et al. 2003; Postma et al. 2004; Rooijakkers et al. 2005; van Wamel et al. 2006; Thammavongsa et al. 2015; van Alen et al. 2018). They are temperate prophages that primarily integrate into the *hlb* gene of *S. aureus*. While ϕSa3 prophages are present in 88% of the human-associated CC398 isolates in our collection, we find that this is not a consequence of a stable association between CC398 and one prophage. Nucleotide diversity within genes shared across ϕSa3 prophages (Figure 4, Supplementary File 3), suggest at least seven (but likely more) acquisitions of an ϕSa3 prophage into human-associated CC398 from outside of CC398. The set of functional genes carried by these elements indicate that the elements within human-associated CC398 include types C (n=285), B (n=111), E (n=35), A (n=4) and D (n=4), and those carried by livestock-associated CC398 isolates include types B (n=84), E (n=8) and A (n=5) (van Wamel et al. 2006; van Alen et al. 2018).

**Figure 4.**
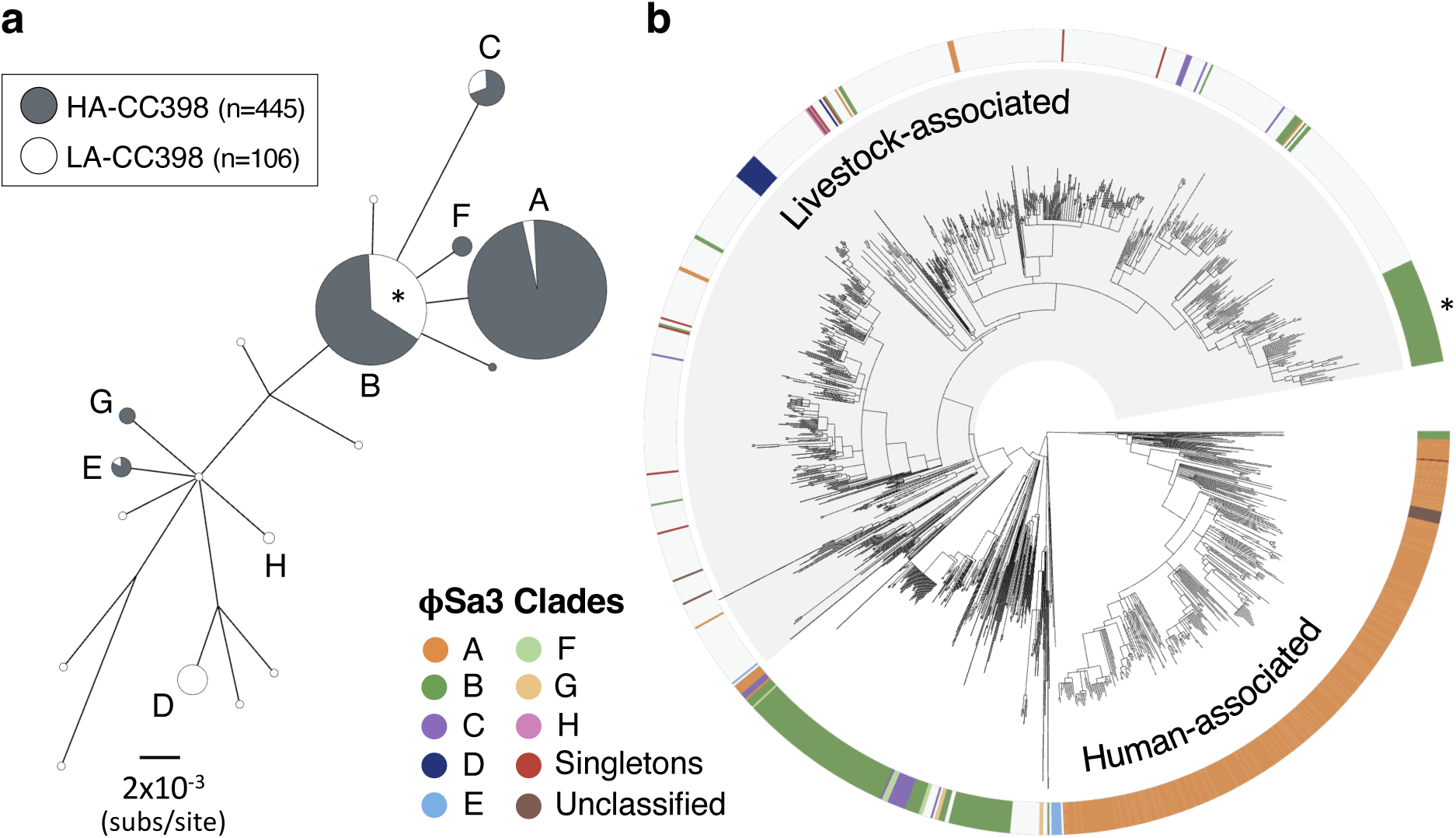
ϕSa3 prophages have been lost and acquired multiple times in both human-associated and livestock-associated CC398. (a) A maximum likelihood phylogeny based on 12 genes shared across the ϕSa3 prophages in our collection, with both low-support nodes (<70% bootstrap support) and branches <0.0018 subs/site collapsed. The latter cut-off is a conservative estimate of the maximum distance that could reflect divergence within CC398. It is the maximum pairwise distance between isolates carrying ϕSa3 prophages across 1,000 estimates from random samples of a core gene alignment of the same number of sites as is in our ϕSa3 prophage alignment. Node size correlates with the number of elements on a log-scale. Elements carried by isolates from human-associated CC398 (grey) and livestock-associated CC398 (white) isolates are indicated, and nodes that include multiple elements labelled (A-E). (b) These clades annotated on the CC398 phylogeny as an external ring. The element carried by the poultry-associated subclade of livestock-associated CC398 is indicated by *.

While ϕSa3 prophages are rare in livestock-associated CC398 (68/704 isolates), diversity within these elements indicate at least 15 (but likely more) acquisitions of these MGEs by livestock-associated CC398. The majority of these elements (69%) do not share a recent common ancestor with those in human-associated CC398. ϕSa3 prophages present in livestock-associated CC398 generally show evidence of recent acquisition, with a notable exception being a previously described poultry-associated sub-clade (n=51) (Price et al. 2012; Larsen et al. 2016b; Pérez-Moreno et al. 2017; Tang et al. 2017). In our collection, 39% of the isolates in this clade are from poultry (compared to 4% across the rest of the livestock-associated clade). Low nucleotide diversity within the type B ϕSa3 consistently present in isolates within this clade suggests that it has been maintained since its acquisition approximately 21 years ago (95% CI: 1997-2001; Figure 1b and Figure 4-figure supplement 1).

### Distinct route to multi-drug resistance in livestock-associated CC398

Livestock-associated CC398 is frequently multi-drug resistant. We see evidence of this in our data set where 81% of livestock-associated CC398 isolates carry one or more resistance genes for antibiotic classes other than tetracyclines and β-lactams (Figure 5-figure supplement 1, Figure 5-figure supplement 2, Supplementary File 7). 67% of livestock-associated CC398 isolates have genes associated with trimethoprim resistance (*dfrA, dfrK* or *dfrG*), 42% have genes associated with macrolide resistance (*ermA, ermB, ermC* or *ermT*), and 26% have genes associated with aminoglycoside resistance (*aadA, aphA* or *aphD*). Not only are resistance genes less common in human-associated CC398 (only 20% of isolates carry tetracycline resistance genes, and 9% carry trimethoprim resistance genes), they also differ in their relative frequencies. In particular, human-associated CC398 isolates more commonly carry genes associated with macrolide resistance (91%).

### Spillover of livestock-associated CC398 into humans is associated with the acquisition of ϕSa3 prophages, but not a loss of resistance genes

ϕSa3 prophages are more common in human isolates (23%) than in livestock or companion animal isolates (11%) in our collection of livestock-associated CC398. To determine the significance of this association, we identified 70 phylogenetically independent clades that include isolates from both human and livestock or companion animal hosts (Figure 5-figure supplement 3). Comparisons across these groups revealed that isolates from humans are consistently more likely to carry an ϕSa3 prophage (McNemar’s Chi-squared test: *p* = 1.50 × 10^−3^; Figure 5).

**Figure 5.**
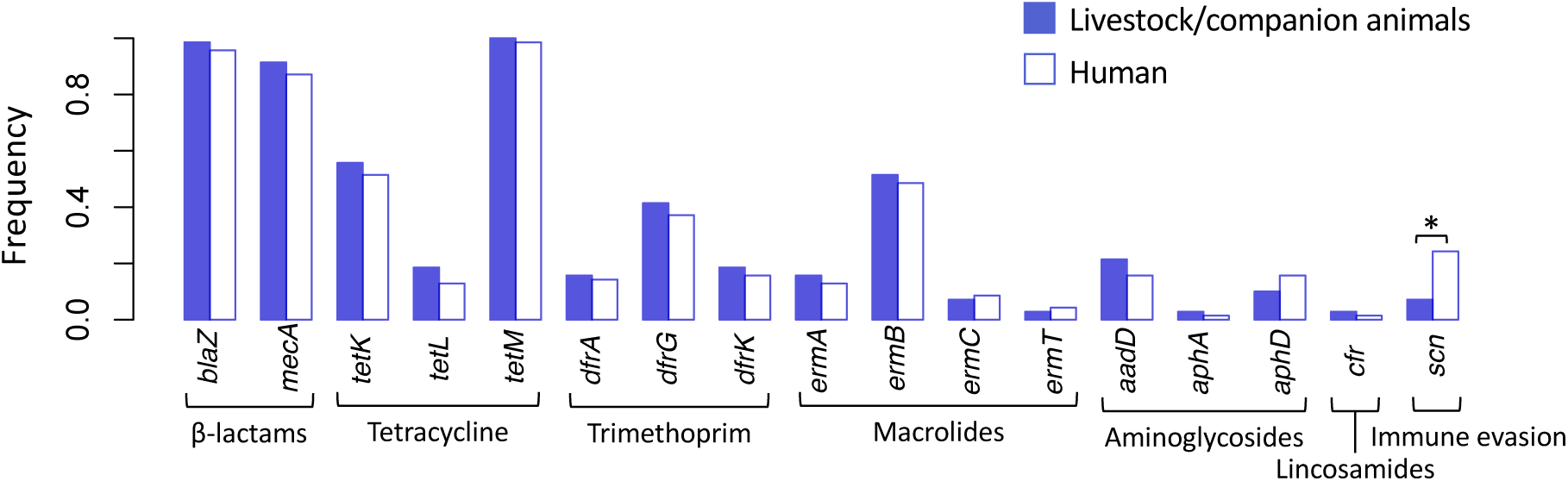
Spillover of livestock-associated CC398 into humans is associated with acquisition of human immune-evasion genes. 70 phylogenetically independent clades that include isolates from both humans and other species were identified within the livestock-associated clade. The plot shows the frequency with which these genes were identified within isolates from humans (right, empty bars) and non-human species (left, filled bars) in these groups. An asterisk indicates a significant difference based on McNemar’s Chi-squared test (*p* = 1.50 × 10^-3^). No resistance genes differed significantly in their frequency across the human and non-human hosts (*p* > 0.1). The *scn* gene is always present in the human immune evasion cluster carried by ϕSa3 prophages, and therefore represents the presence of this element.

ϕSa3 prophages are also less common in isolates from pigs than from other non-human species (only 3% of pig isolates carry one). In 42/70 of our phylogenetically independent groups, the only non-human species was a pig, and none of the pig isolates from these groups carried an ϕSa3 prophage (while 14% of the human isolates did). In the remaining 28 groups (that included isolates from cows, horses and poultry), ϕSa3 prophages were observed in non-human hosts in 17% of groups, but still at a higher frequency in humans (39% of groups) (McNemar’s Chi-squared test: *p* = 0.04).

In contrast, we found no evidence that the spillover of livestock-associated CC398 into humans is associated with the loss (or gain) of individual antibiotic resistance genes (Figure 5). This contrasts with the conclusions of a previous study that suggested that the resistance genes carried by livestock-associated CC398 are likely to be lost in human hosts (Sieber et al. 2019). We find that no resistance gene was significantly more or less common in humans than in livestock species (McNemar’s Chi-squared tests: *p* > 0.1), nor was there a consistent shift in the overall number of resistance genes (there were more resistance genes in isolates from human hosts in 22/70 pairs and more in livestock hosts in 34/70 pairs).

## Discussion

We have characterised the evolutionary dynamics of the three classes of MGEs that show the greatest changes in frequency across human-and livestock-associated CC398: the Tn*916* transposon and the SCC*mec* element, which are both common in livestock-associated CC398, and the ϕSa3 prophage, which is common in human-associated CC398. Despite a consistency in the relative frequencies of these elements across CC398, leading to their strong association with the transition to livestock, these three elements show a broad spectrum of dynamics. The Tn*916* transposon carrying *tetM* shows evidence of stable and consistent vertical transmission in livestock-associated CC398 and absence from human-associated CC398. The type Vc SCC*mec*, carrying not only *mecA*, but also *tetK* and *czrC*, has also been stably maintained by several lineages of livestock-associated CC398, but by a combination of vertical and horizontal transmission, and with occasional replacement with type IV elements. While type V SCC*mec* elements are also present in human-associated CC398, there is little evidence of their longer-term maintenance. Finally, while ϕSa3 prophages carrying a human immune-evasion gene cluster are rare in livestock-associated CC398 and common in human-associated CC398, there have been frequent gains and losses in both groups. These contrasting dynamics may reflect variation in the selective benefits and costs of the carriage of these MGEs by CC398, their availability in the environments encountered by CC398, and their mechanisms of transfer.

The three classes of MGEs whose dynamics we describe all employ different mechanisms of horizontal transfer, and this may both influence their intrinsic stability in the CC398 genome and their availability for acquisition from outside of CC398. Experimental studies have found that different types of MGEs vary in their rates of transfer between bacterial cells. In particular, in vitro rates of transfer of phage have been found to be several orders of magnitude higher than of transposons (Humphrey et al. 2021). A 16-day in vivo study of MGE dynamics for two strains of CC398 also observed variation in the mobility of MGEs: the Tn*916* transposon, the type V SCC*mec* and an ϕSa3 prophage were all stably maintained and not horizontally transferred, while other prophages and plasmids were both gained and lost by daughter cells (McCarthy et al. 2014). While variation in the intrinsic stability of MGEs in their host will influence their long-term dynamics, other factors may dominate. For instance, the long-term stability of Tn*916* within CC398 could emerge from a broad spectrum of short-term dynamics. It could result from a very low rate of loss, or frequent loss combined with a high selective cost of loss and a low probability of reacquisition from sources other than very closely related cells. Therefore, the long-term dynamics we describe provide a unique insight into the relationship between these MGEs and CC398.

We find that a Tn*916* transposon that carries *tetM* was acquired once by CC398 and this coincided with its origin in European livestock in 1964 (95% CI: 1957-1970). Our observation of the loss of Tn*916* on terminal branches of the phylogeny suggests that while losses do occur, lineages that lose Tn*916* are either rapidly outcompeted by those that have maintained it, or they rapidly reacquire it from close relatives. Antibiotics, including tetracyclines, were first licensed for use as growth promoters in livestock in European countries in the 1950s, and were in common use by the end of that decade (Lowbury et al. 1958). While the use of antibiotics as growth promoters was banned by the European Union in 2006, tetracyclines remain the most commonly used antimicrobial class in livestock farming (WHO 2015; DANMAP 2019). Carriage of Tn*916* is therefore likely to have been associated with a strong selective benefit for livestock-associated CC398 ever since its emergence. However, as CC398’s exposure to tetracyclines will be intermittent, the long-term stability of Tn*916* is also likely to reflect a low selective cost in the absence of treatment or a barrier to reacquisition following loss. Tn*916*-like elements are found across several bacterial genera (Clewell et al. 1995; Roberts and Mullany 2009), including other opportunistic pathogens in the respiratory microbiome of pigs (Holden et al. 2009; Hoa et al. 2011) and other *S. aureus* CCs (de Vries et al. 2009). This suggests that the stability of Tn*916* in CC398 is not due to the rarity of the element in the environments encountered by CC398, however it may be due to other barriers to successful transfer.

Previous studies have found evidence that the regulatory system of Tn*916* promotes its maintenance in a host through ensuring both that excision only occurs in the presence of tetracycline (or other transcription-limiting cell stress) and that any selective burden in the absence of treatment is minimised (Roberts and Mullany 2009). Nevertheless, Tn*916* has been found to have a much more dynamic association with lineages of other bacterial species (e.g. *Streptococcus pneumoniae* (D’Aeth et al. 2021)). The fitness costs of MGEs can be host (and even insertion locus) specific and can also be mitigated over time (Starikova et al. 2013; Durão et al. 2018), and therefore the stability of Tn*916* in livestock-associated CC398 might reflect a low cost that is specific to this lineage. A combination of high selective benefit, low selective cost and inaccessibility for reacquisition following loss could explain the remarkable stability of Tn*916* in livestock-associated CC398, and may in part explain the success of this lineage in livestock.

Our results indicate that the acquisition of an SCC*mec* occurred later in the expansion of livestock-associated CC398. The most common SCC*mec* in livestock-associated CC398, type Vc, carries *tetK* and *czrC* in addition to *mecA*. These additional resistance genes might be highly advantageous in livestock, and particularly in pigs. While all livestock-associated CC398 carry *tetM*, there is evidence that carrying *tetK* in addition to *tetM* is associated with increased fitness during exposure to sublethal concentrations of tetracycline (Larsen et al. 2016a). *czrC* is associated with heavy metal resistance (Cavaco et al. 2010), which is likely to be beneficial in the context of the common supplementation of animal feed with zinc oxide, which in pigs is commonly used to prevent diarrhoea in weaners (Nielsen et al. 2021). Additionally, *mecA* is likely to be beneficial because of the common use of beta-lactams in livestock farming, including third generation cephalosporins (Sjölund et al. 2016; Lekagul et al. 2019). The size of the type Vc element, combined with the replacements and truncations we observe suggests that it may come with a selective cost. In addition, SCC*mec* type Vc appears to be rare (at least in *S. aureus*). The element has been found in other staphylococci (*S. cohhii* in Vervet Monkeys (Hoefer et al. 2021)), but has not been reported in other *S. aureus* CCs.

Loss of the type Vc SCC*mec* on internal branches within livestock-associated CC398 is associated with replacement with a type IV SCC*mec* that only carries *mecA*. While we find evidence of only a single acquisition of the type Vc SCC*mec* by livestock-associated CC398, we find evidence of at least four acquisitions of type IV SCC*mec*. The more recent dates of acquisition, and an apparent association with livestock species other than pigs, might reflect a difference in selective pressures across different livestock species, or a loss of the type Vc element during transmission between livestock populations. The long-term maintenance of the type Vc SCC*mec* is consistent with a high selective benefit (particularly in pigs where this element is most common). However, its repeated loss and replacement with type IV elements is consistent with a low cost of this replacement, at least in certain contexts (perhaps in other livestock hosts), and might also reflect the rarity of the type V element relative to the type IV.

While the loss of the ϕSa3 prophage that carries a human immune-evasion gene cluster is associated with the transition to livestock, neither the loss nor the gain of these elements is likely to be a substantial hurdle for the adaptation of CC398 to human or non-human hosts. Their ubiquity in human-associated CC398 and frequent acquisition following transmission of livestock-associated CC398 to humans (consistent with Sieber et al. 2019), is consistent with a strong benefit in the human host (Rohmer and Wolz 2021). On the other hand, as we find that these elements are frequently lost and generally absent in other host species, these elements may carry a selective burden outside of the human host. The diversity of ϕSa3 prophages in CC398 suggests they form a large pool of elements within human hosts (van Alen et al. 2018), likely reflecting their ubiquity in human carriage populations of *S. aureus* (Rohmer and Wolz 2021) and capacity to transfer between *S. aureus* lineages.

We observe one clear exception to the pattern of recent acquisitions of ϕSa3 in livestock-associated CC398 in response to spillover events: the acquisition of an ϕSa3 prophage at the base of a poultry-associated sub-clade of livestock-associated CC398 (Price et al. 2012; Larsen et al. 2016b; Pérez-Moreno et al. 2017; Tang et al. 2017). This clade was first described as a hybrid LA-MRSA CC9/CC398 lineage (Price et al. 2012), and was subsequently investigated as a lineage associated with human disease (Larsen et al. 2016b). The maintenance of an ϕSa3 in this lineage may reflect more frequent transmission via a human host, or an adaptation to poultry (a recent study suggested that ϕSa3 prophages might aid immune evasion in species other than humans (Jung et al. 2017)). Either way, this might make this lineage a greater immediate threat to public health.

While livestock-associated CC398 is found across a broad range of livestock species, it is most commonly associated with pigs. The dynamics of the SCC*mec* and ϕSa3 prophages that we have identified are both consistent with livestock-associated CC398 originating in pig farms and later spreading to other livestock species. The lower frequency of the type Vc SCC*mec* in species other than pigs could reflect random loss during transmission bottlenecks, or a reduced benefit of either *tetK* or *czrC* in these species. Similarly, the higher frequency of ϕSa3 prophages in other species might reflect an increased benefit or reduced cost, or more recent transmission via human hosts.

Our results reveal that LA-MRSA CC398 is a stably antibiotic-resistant pathogen that is capable of dynamic readaptation to humans. We find that Tn*916* and SCC*mec* are both stably maintained in livestock-associated CC398, across different livestock species and countries, and that neither of these MGEs (or other antibiotic resistance genes) tend to be lost when livestock-associated CC398 is transmitted to humans. This suggests that these MGEs are associated with a low selective cost in both livestock and human hosts, and therefore across variable levels and types of antibiotic exposure. The stability of these two MGEs, combined with the capacity of livestock-associated CC398 to rapidly acquire the ϕSa3 prophage, underlines the threat posed by LA-MRSA public health. While SCC*mec* is less stably maintained than Tn*916*, our identification of several independent acquisitions of type IV SCC*mec* suggests both a strong selective benefit, not contingent on the carriage of *tetK* and *czrC*, and an availability of type IV elements. These dynamics predict that the impact of gradual reductions in antibiotic consumption on LA-MRSA are likely to be slow to be realised and that the forthcoming EU ban on medical zinc supplementation in pig feed may have a limited impact on LA-MRSA (European Medicines Agency 2017, 2020; DANMAP 2019). Further work is, however, required to understand the factors that underlie the acquisition and maintenance of resistance genes within LA-MRSA, and how they differ from human-associated MRSA lineages.

## Methods

### Data collection

All available genome sequence data relating to *S. aureus* CC398 was downloaded from public databases (www.ncbi.nlm.nih.gov/sra and www.ebi.ac.uk/ena; accessed 2019), with metadata in some cases obtained by request (Supplementary File 1). We additionally sequenced five isolates sampled from UK pig farms in 2018. All publicly available complete *S. aureus* genomes assemblies (www.ncbi.nlm.nih.gov; accessed 2019) were MLST typed using Pathogenwatch (https://pathogen.watch), and the 43 genomes identified as CC398 were added to our collection. After exclusion of low-quality assemblies and isolates mischaracterised as CC398, this led to a collection of 1,180 genomes.

### Genomic library preparation and sequencing

For sequencing of the five UK pig farm isolates, genomic DNA was extracted from overnight cultures grown in TSB at 37°C with 200rpm shaking using the MasterPure Gram Positive DNA Purification Kit (Cambio, UK). Illumina library preparation and Hi-Seq sequencing were carried out as previously described (Harrison et al. 2013).

### Genome assembly

We used sequence data from all isolates to generate *de novo* assemblies with *Spades* v.3.12.0 (Bankevich et al. 2012). We removed adapters and low quality reads with *Cutadapt* v1.16 (Martin 2011) and *Sickle* v1.33(Joshi and Fass 2011), and screened for contamination using *FastQ Screen* (Wingett and Andrews 2018). Optimal k-mers were identified based on average read lengths for each genome. All assemblies were evaluated using *QUAST* v.5.0.1(Gurevich et al. 2013) and we mapped reads back to *de novo* assemblies to investigate polymorphism (indicative of mixed cultures) using *Bowtie2* v1.2.2 (Langmead and Salzberg 2012). Low quality genome assemblies were excluded from further analysis (i.e., N50 <10,000, contigs smaller than 1kb contributing to >15% of the total assembly length, total assembly length outside of the median sequence length +/- one standard deviation, or >1,500 polymorphic sites). We identified genomes mischaracterised as CC398 via two approaches and excluded them from further analysis. First, we identified sequence types (STs) with *MLST-check* (Page et al. 2016) and grouped into CCs using the *eBURST* algorithm with a single locus variant (Francisco et al. 2009). Second, we constructed a neighbour-joining tree based on a concatenated alignment of MLST genes (*arcC, aroE, glpF, gmk, pta, tpi* and *yqiL*) for our collection and 13 additional reference genomes from other CCs, using the *ape* package in *R* and a K80 substitution model (Paradis et al. 2004).

We generated reference-mapped assemblies with *Bowtie2* using the reference genome S0385 (*GenBank* accession no. AM990992). For reference genomes, we generated artificial FASTQ files with *ArtificialFastqGenerator* (Frampton and Houlston 2012). Average coverage and number of missing sites in these assemblies were used as an additional quality control measure; genomes with average coverage <50x or with >10% missing sites were excluded.

We identified recombination in the reference-mapped alignment using both *Gubbins* v2.3.1(Croucher et al. 2015) and *ClonalFrame* (Didelot and Wilson 2015). We masked all the recombinant sites identified from our alignment. We additionally masked a region of ∼123 kb that was identified as horizontally acquired from an ST9 donor in a previous study (Price et al. 2012).

### Genome annotation and identification of homologous genes

We annotated *de novo* assemblies with *Prokka* v2.8.2 (Seemann 2014) and identified orthologous genes with *Roary* (Page et al. 2015) using recommended parameter values. We created a core gene alignment with *Roary* and identified recombinant sites using *Gubbins*. We identified antibiotic resistance genes using the *Pathogenwatch* AMR prediction module (Wellcome Sanger Institute), which uses *BLASTn* (Altschul et al. 1990) with a cut-off of 75% coverage and 80-90% percent identity threshold (depending on the gene) against a *S. aureus* AMR database.

### Phylogenetic analyses

We carried out phylogenetic reconstruction for the reference-mapped alignment with *RAxML* v8.2.4 using the *GTR*+*Γ* model and 1,000 bootstraps(Stamatakis 2014). Sites where >0.1% of genomes showed evidence of recombination or had missing data were excluded from the analysis. We constructed dated phylogenies using *BEAST* v1.10 with a *HKY*+*Γ* model, a strict molecular clock, and constant population size coalescent prior, from the coding regions of the reference-mapped alignment (Drummond et al. 2012). We fit separate substitution models and molecular clocks to 1st/2nd and 3rd codon positions to reflect differences in selective constraint. We constructed phylogenies for three random subsamples of 250 isolates (200 from livestock-associated CC398 and 50 from human-associated CC398). Subsamples were non-overlapping except for 30 genomes representing the most divergent lineages within the livestock-associated clade, to ensure a consistent description of the origin of this clade. We constructed additional phylogenies from subsamples that included only isolates from the LA-clade to establish that consistent rate estimates were returned over different evolutionary depths. We investigated temporal signal in each data set through a regression of root-to-tip distance against sampling date, and a permutation test of dates over tips (with clustering used to correct for any confounding of temporal and genetic structures) (Murray et al. 2016; Rambaut et al. 2016). Trees are visualized and annotated using *ITOL* (Letunic and Bork 2021).

### Comparative analyses of mobile genetic elements

Genes associated with the transition to livestock were identified through comparing frequencies of homologous genes identified with *Roary* across human- and livestock-associated CC398. We investigated the association between these genes and MGEs through analysis of (i) physical locations within our *de novo* assemblies and the reference genomes, (ii) correlations in their presence/absence across our collection, and (iii) comparison with descriptions in the literature and in online databases of particular MGEs.

We confirmed the identity of the Tn*916* transposon by comparison with published descriptions and publicly available annotated sequences. SCC*mec* elements were initially categorised into types using a *BLASTn* search of all representative SCC*mec* types from the *SCCmecFinder* database in our *de novo* assemblies (Supplementary File 5) (Hanssen and Ericson Sollid 2006). ϕSa3 prophages were initially identified and categorised through identification of functional genes associated with the human immune evasion gene cluster. To further characterise diversity within these elements we identified genes associated with each element across our collection. We constructed alignments of genes within MGEs using *Clustal Omega* v1.2.3 (Sievers and Higgins 2018) and checked for misalignment by eye.

We analysed variation in both gene content and nucleotide diversity within shared genes for each MGE. We estimated pairwise nucleotide distances in concatenated alignments of shared genes for each type of element using the *ape* package in *R* (Paradis et al. 2004), and constructed maximum likelihood trees using *RaxML* and minimum spanning trees using *GrapeTree* (Zhou et al. 2018) to investigate co-phylogeny. We generated confidence intervals for our estimates of mean pairwise nucleotide distances within MGEs by re-estimating the mean distance from 1,000 bootstrapped samples of sites. We compared this to sites in the core genome by sampling the same number of sites as were in the MGE alignment from a concatenated alignment of core genes (generated by *Roary*). For SCC*mec* we used ancestral state reconstruction in *BEAST* to infer the evolutionary dynamics of these elements and date their origins within CC398. This involved fitting a discrete traits model to the posterior distributions of trees, with each state representing a version of the element that had been independently acquired by CC398. We used a strict clock model that allowed for asymmetric rates of transitions between states, but we found that the results were robust to use of a symmetric model or a relaxed clock

### Phylogenetically independent groups

To test the association between spillover into the human host and the presence of ϕSa3 prophages and antibiotic-resistance genes, we identified 70 phylogenetically independent clades of isolates that were sampled from both human and non-human hosts in the livestock-associated clade. We classed a gene as present in a host if it was observed in any of the isolates from that host in that clade.

## Ethics approval and consent to participate

Not applicable.

## Consent for publication

Not applicable.

## Availability of data and materials

The Illumina sequences generated in this study have been deposited in the NCBI short read archive under the accession numbers ERR3524650, ERR3524328, ERR3524354, ERR3524446 and ERR3524562. All other sequences used in this study are publicly available and their origins are described in Supplementary File 1.

## Acknowledgements

MM was funded by the Medical Research Council, co-funded by the Raymond and Beverly Sackler Fund. GGRM and LAW were supported by a Sir Henry Dale Fellowship jointly funded by the Wellcome Trust and the Royal Society (109385/Z/15/Z). GGRM was also supported by a ZELS BBSRC award (BB/L018934/1) and a Research Fellowship at Newnham College.

## Supplementary Figure Legends

**Figure 1-figure supplement 1. The temporal, host-species and geographic distribution of our collection of CC398 isolates**. (a) Phylogeny of CC398 with rings showing the host species and countries of origin for each isolate (groups with n<10 not shown). The blue outline indicates the livestock-associated clade, and the red outline indicates the 5 most recent isolates in our collection (sampled in 2018), from pigs on UK farms. (b), (c) and (d) show the variation in sampling date, host species and country across the livestock-associated (blue, lower) and human-associated (red, upper) clades.

**Figure 1-figure supplement 2. Different outgroups consistently identify the root of CC398 within human-associated CC398**. We constructed maximum likelihood phylogenies using a reference-mapped alignment of CC398 combined with four outgroups from sequence types (STs) 291, 30, 97 and 5; and using a core genome alignment of our *de novo* assemblies and a midpoint rooting. The outgroups we used covered a range of distances from the base of CC398: ST291 (ERR2729529) is∼0.005 subs/site, ST30 (ERS1420125) is∼0.012 subs/site, and ST97 and ST5 (ERR2729579 and SRS613151) are∼0.015 subs/site. The reference-mapped phylogenies that were rooted using ST291, ST30 and ST97, and the midpoint-rooted core genome phylogeny all showed a consistent root, which is shown in the figure (and indicated by 1). A different root was obtained when we used the outgroup of ST5 (location indicated by 2). This root is on a neighbouring branch to the root found in the 4 other reconstructions, and results in CC398 being rooted on the branch leading to a single isolate (ZTA09_03734_9HSA). Livestock-associated CC398 is indicated by a blue box with grey shading.

**Figure 1-figure supplement 3. A consistent estimate of the age of the livestock-associated clade**. Results of BEAST dating analyses estimating (a) the origin of a shallower subclade within the livestock-associated clade, (b) the origin of the entire livestock-associated clade, and (c) the origin of CC398. (a) The figure shows a schematic representation of the CC398 phylogeny indicating the nodes of interest (A-C), and our sampling strategy. We randomly sampled our data set three times to generate samples of 250 isolates (200 from the livestock-associated clade and 50 from the human-associated clade). Samples overlapped by only 30 isolates that represent the most divergent lineages of the livestock-associated clade, to ensure a consistent description of the most recent common ancestor. Each sample had the same range of sampling dates (1993-2018). (a) also describes the results of a regression of root-to-tip distance against sampling date (correlation coefficient and estimate of the date of the most recent common ancestor) for each sample (1-3), and subsamples that include isolates from (A) only the main livestock-associated clade, (B) only the livestock-associated clade, and (C) the entire sample. As we observed stronger temporal signal for A and B, than for C, we estimated dated trees using BEAST for each of these 9 subsampled data sets. We observed consistent estimates of evolutionary rate across all these analyses (b). Rates at 1st/2nd codon positions are shown as circular points, and at 3rd codon positions as square points). These analyses also returned broadly consistent estimates of dates (c). Although we found that estimates from subsamples that included outgroups of the node being dated returned more precise and marginally more recent estimates of age, likely due to more information about the location of the root.

**Figure 1-figure supplement 4. Evidence of temporal signal is present across in our subsampled datasets, but is stronger when isolates from the human-associated group are excluded**. Regressions of root-to-tip distance against sampling date for each of our data sets, rooted to minimise residual mean squares. (a)-(c) show the results for samples 1, 2 and 3 for clade a (described in Figure 1-figure supplement 3), (d)-(f) the results for samples 1, 2 and 3 for clade b, and (g)-(i) samples 1, 2 and 3 for clade c. For all data sets a randomisation test indicated that these correlations were unlikely to have arisen by change (*p*<0.01). Correlation coefficients (r) and estimates of the time of the most recent common ancestor (tMRCA) based on the regression are shown for each data set.

**Figure 1-figure supplement 5. Livestock-associated and human-associated CC398 have divergent accessory genomes, and genes whose presence/absence most clearly distinguish these groups (except one) are associated with a Tn*916* transposon, SCC*mec* and ϕSa3 prophages**. (a) A plot of the 1st and 2nd principal components of accessory genome content, with isolates from human-associated CC398 in red and isolates from livestock-associated CC398 in blue. This was constructed using the package *adegenet* in *R* [1]. (b) Comparison of gene frequencies across the human-associated and livestock-associated groups. Genes present in <20% of human-associated CC398 and >80% of livestock-associated CC398 and genes presence in >80% of human-associated CC398 and <20% livestock-associated CC398 are highlighted, with genes associated with the Tn*916* transposon shown in purple (all are overlapping as they have identical frequencies), genes associated with SCC*mec* shown in turquoise, and genes associated with ϕSa3 prophages shown in blue. The one gene that distinguishes livestock-associated CC398 from human-associated CC398 that isn’t associated with one of these three MGEs is shown as a red circle (*tatC*).

**Figure 2-figure supplement 1. Evidence of repeated excision of Tn*916* transposon in the livestock-associated clade**. (a) Purple points indicate which 5 isolates lack the Tn*916* transposon in the livestock-associated clade. (b) Top: A gene map showing the Tn*916* transposon from the 1_1429 reference strain and the two flanking genes (dark grey). Bottom: The integration site in the 62951 strain within which the transposon is absent. The percentage nucleotide identity between the two flanking genes is indicated below the bars connecting the top and bottom gene maps.

**Figure 2-figure supplement 2. Type V and IV SCC*mec* elements identified in CC398 through a *BLASTn* search or representative types**. Initial identification of SCC*mec* types was carried out by *BLASTn* search of all SCC*mec* reference sequences on the *SCCmecFinder* extended database. As the SCC*mec* element in our dataset was commonly separated onto multiple contigs, we found that contiguous hits of the entire element were rare. Therefore, to determine the presence of a particular SCC*mec* type we considered the combined length of high-identity hits (hits that have >95% nucleotide identity and are ≥5% of the length of the element). We categorised elements into SCC*mec* types based on the overall length of the matched region, and described strains with best match lengths of <50% as unknown. The CC398 phylogeny is annotated with the results of this analysis. The innermost ring shows the presence/absence of *mecA*, and outer rings show the percentage length match for each type/sub-type (Supplementary File 7).

**Figure 3-figure supplement 1. Most of the shorter versions of the type Vc SCC*mec* element in CC398 can be attributed to deletion events**. Of the 540 isolates that carry a type V SCC*mec* in CC398, 204 have >2 genes absent. There are 3 common forms of gene absence (B, C and D). (a) Gene maps showing common patterns of gene absence. (b) A minimum spanning tree of a concatenated alignment of the 40 genes associated with the full-length SCC*mec* type V. Points represent groups of elements that differ by a maximum of 1 SNP, with point size correlated with group size on a log-scale, and colours representing categories described in (a). (c) Annotation of the full-length and truncated categories onto the core genome phylogeny. (d) Histograms showing pairwise distances (per shared nucleotide site) between individual elements in each group and the least divergent member of Group A. Histograms are shown on a log(x+1) scale due to the high frequency of low divergence elements. The dashed vertical lines show the average pairwise distance between isolates within the livestock-associated clade based on a core gene alignment, and the dotted lines show 10-times this value. Groups A, B and C are only found in the livestock-associated clade. Due to the low level of diversity in group D, versions from the human-associated and livestock-associated clade are shown together, while for group E they are shown separately (HA/LA). There are 13 isolates from the human-associated clade in group E. Of these 13 isolates, only 3 show sufficiently low divergence to be consistent with a recent common ancestor with the majority of elements from Group A, within CC398 (for these three elements divergence is <1.7 ×10^− 4^ /site). The three isolates carrying these elements all fall within the recent Danish hospital outbreak clade, and show 100% identity with the majority of the elements within this clade (that fall within group D).

**Figure 3-figure supplement 2. There have been at least four independent acquisitions of type IV SCC*mec* within livestock-associated CC398**. (a) An minimum-spanning tree based on a concatenated alignment of the 12 genes shared across all type IV SCC*mec* (n=148 isolates). Clades are distinguished based on nucleotide distances within these shared genes and by differences in the genes present in these elements. Clade A is IVa(1), Clade B is IVc, Clade C is IVa(2), and Clade D is IVa(3). IVa(2) and IVa(3) only differ by one SNP in the alignment of 12 shared genes, but they have different gene contents, suggesting divergent origins. (b) Clades of type IV elements mapped onto the core genome phylogeny.

**Figure 3-figure supplement 3. Tn*916* was acquired before the current complement of SCC*mec* elements in livestock-associated CC398, SCC*mec* type V was acquired before SCC*mec* type IV, and SCC*mec* type V was acquired by livestock-associated CC398 before human-associated CC398**. Date estimates based on ancestral state reconstructions with BEAST over dated phylogenies inferred from three subsampled data sets. (A) A dated phylogeny inferred from one subsample (no. 1) with branches coloured by inferred SCC*mec* element at their descendent node (>95% posterior support), and earliest nodes (>95% posterior support) for each element labelled. (B) Date estimates for the earliest nodes at which each SCC*mec* type is inferred to be present (>95% posterior support) across the three data sets. Points are median values and bars are 95% confidence intervals. Estimates are shown for three versions of SCC*mec* type IVa that were distinguished based on the divergence within this element (see Figure 3-figure supplement 3). Only one estimate is provided for SCC*mec*-IVa(3) because it is only observed in one of the subsampled data sets.

**Figure 4-figure supplement 1. Maintenance of a ϕSa3 prophage in a poultry-associated clade of livestock-associated CC398 for around 21 years**. (a) A minimum spanning tree of the type B ϕSa3 prophage present in the poultry-associated clade of livestock-associated CC398. 55 isolates in our collection of CC398 carry a version of this element. We identified 34 genes shared across these 55 versions of the element. After pruning genes with homologues, and difficult to align regions 29/34 genes were used to construct the tree. Points represent groups of identical elements, with point size correlated with group size on a log-scale, and colours representing clades that differ from the basal group by >1 SNP. (b) These clades annotated onto the CC398 phylogeny as an external ring. 51/55 isolates carrying this element fall within the poultry-associated clade and 4 other isolates are from other clades within livestock-associated CC398 from human hosts. Within the avian clade we see little diversity (maximum pairwise distance is 1 SNP). The elements from outside of this clade show greater divergence, but this could still represent divergence within livestock-associated CC398.

**Figure 5-figure supplement 1. Patterns in the presence/absence of antibiotic resistance genes vary across livestock-associated CC398 and human-associated CC398 in addition to the three genes associated with Tn*916* and SCC*mec***. Variation in presence of antibiotic resistance genes across CC398 mapped onto the core genome phylogeny. All genes that are present above a threshold of 5% are represented, with colours representing different antibiotic classes.

**Figure 5-figure supplement 2. Differences in antibiotic resistance gene frequencies across human-and livestock-associated CC398**. (a) Comparison of the frequencies of resistance genes across clades, with the livestock-associated group shown in the filled bar on the left and the human-associated group the empty bar on the right. (b) Comparison of only isolates carrying *mecA* from the two groups. (c) and (d) Venn diagrams that show the relationship between the presence of genes associated with different antibiotic classes in MRSA isolates from both groups.

**Figure 5-figure supplement 3. Phylogeny of the livestock-associated clade showing locations of 70 phylogenetically independent groups of isolates from human (red) and livestock/companion animal (blue) hosts**. The phylogeny is a maximum likelihood phylogeny of the livestock-associated clade, rooted using human-associated CC398, with nodes with <70% bootstrap support collapsed. 70 well-supported independent clades that contain both isolates from human and livestock/companion animal hosts were identified across the livestock-associated clade, and were used to test for differences in the frequency of genes associated with both antibiotic resistance and adaptation to the human host across these two groups. The small number of isolates from wild or domestic animals (other than livestock and horses) were excluded from the analysis.

## Supplementary Table Legends

**Supplementary File 1. Metadata for all isolates used in this study**. Strain names, country of origin, source (host species), year, accession numbers and references for all isolates.

**Supplementary File 2. Genes that most strongly distinguish human-and livestock-associated CC398, and their association with mobile genetic elements**. Our gene identifiers and the gene locations and locus tags in published reference genomes are provided.

**Supplementary File 3. Description of the presence of MGEs and annotation of MGE types and clades**. The presence/absence of genes and MGEs are described by 1/0, and types and clades that are presented in the text are described.

**Supplementary File 4. Description of the genes in the Tn*916* element**. The genes in the Tn*916* element used in our analyses are described in the reference genome 1_1439.

**Supplementary File 5. Reference SCC*mec* elements used in BLAST typing**.

**Supplementary File 6. Description of the genes in the type V SCC*mec* element**. The genes in the SCC*mec* type Vc element used in our analyses are described in the reference genome 12_LA_293.

**Supplementary File 7. AMR genes identified by PathogenWatch**. Gene presence/absence is described by 1/0.

