## Supplementary figures and images for "Stable antibiotic resistance and rapid human adaptation in livestock-associated MRSA"

### Figure1 - Figure Supplement 4

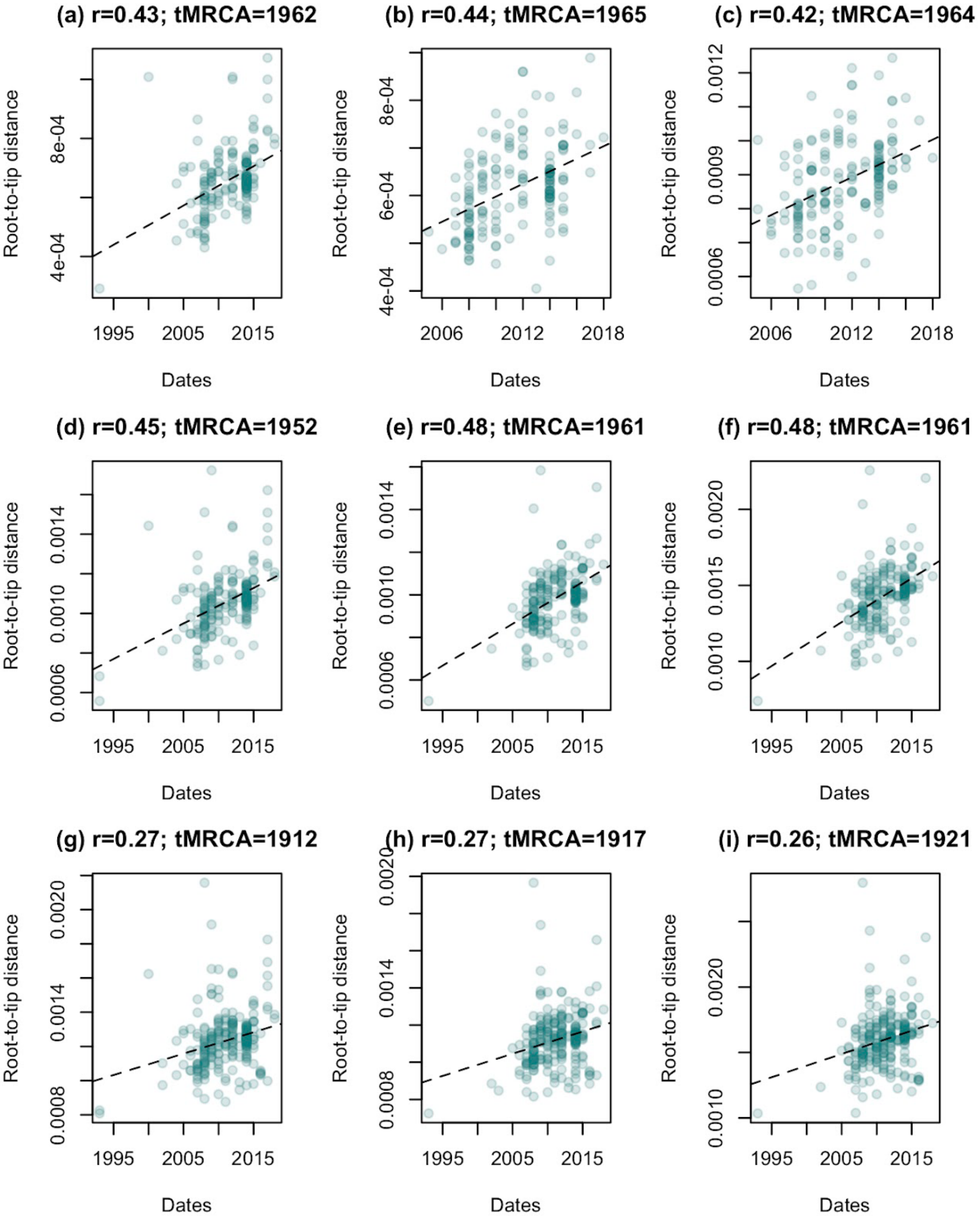

### Figure 1 - Figure Supplement 1

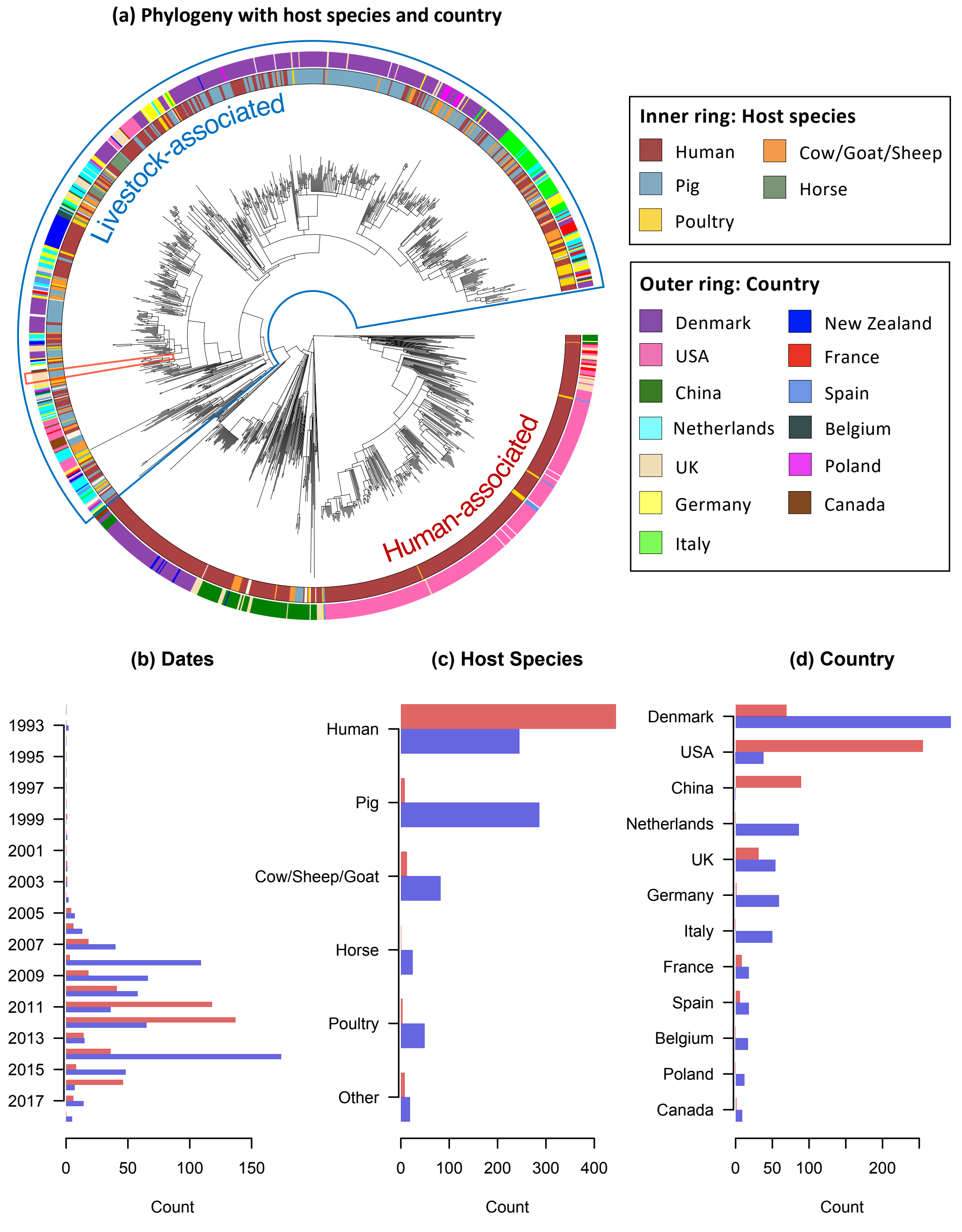

### Figure 1 - Figure Supplement 2

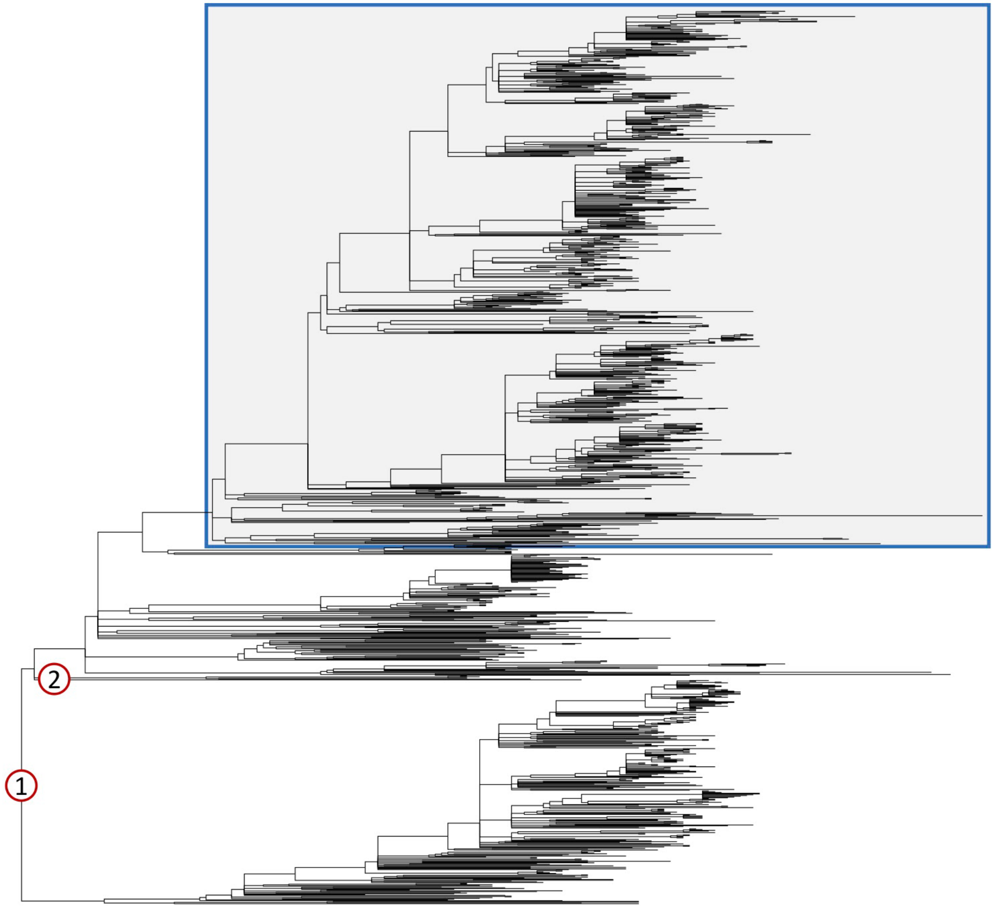


① ST291, ST30, ST97, and midpoint core genome
② ST5

### Figure 1 - Figure Supplement 3

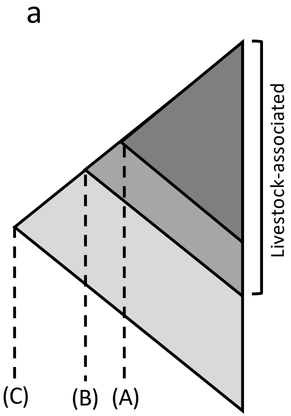


| Clade | Sample | *r* | tMRCA |
|---|---|---|---|
| a | 1 | 0.43 | 1962 |
| | 2 | 0.44 | 1965 |
| | 3 | 0.42 | 1964 |
| b | 1 | 0.45 | 1952 |
| | 2 | 0.48 | 1961 |
| | 3 | 0.48 | 1961 |
| c | 1 | 0.27 | 1912 |
| | 2 | 0.27 | 1917 |
| | 3 | 0.26 | 1921 |

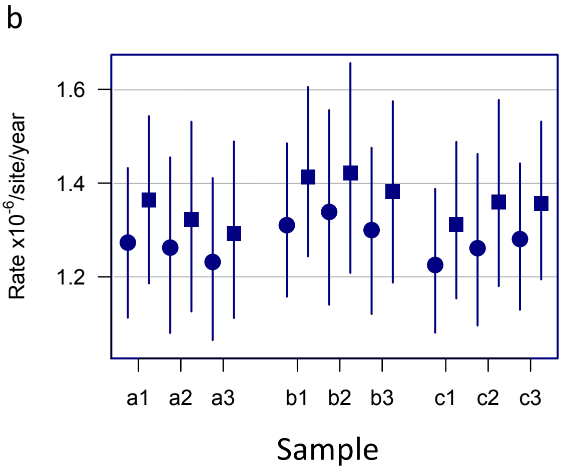


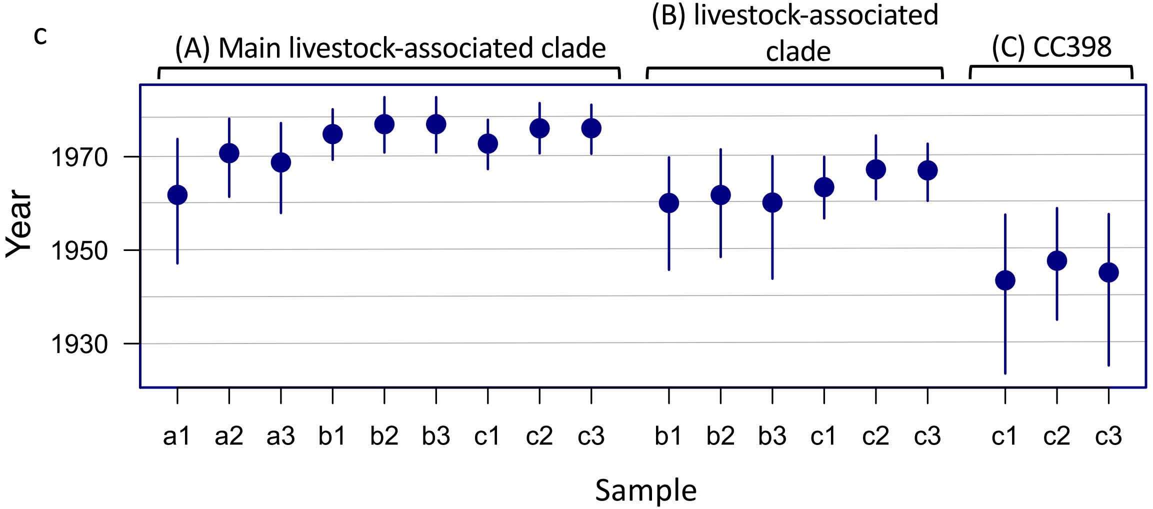

### Figure 1 - Figure Supplement 5

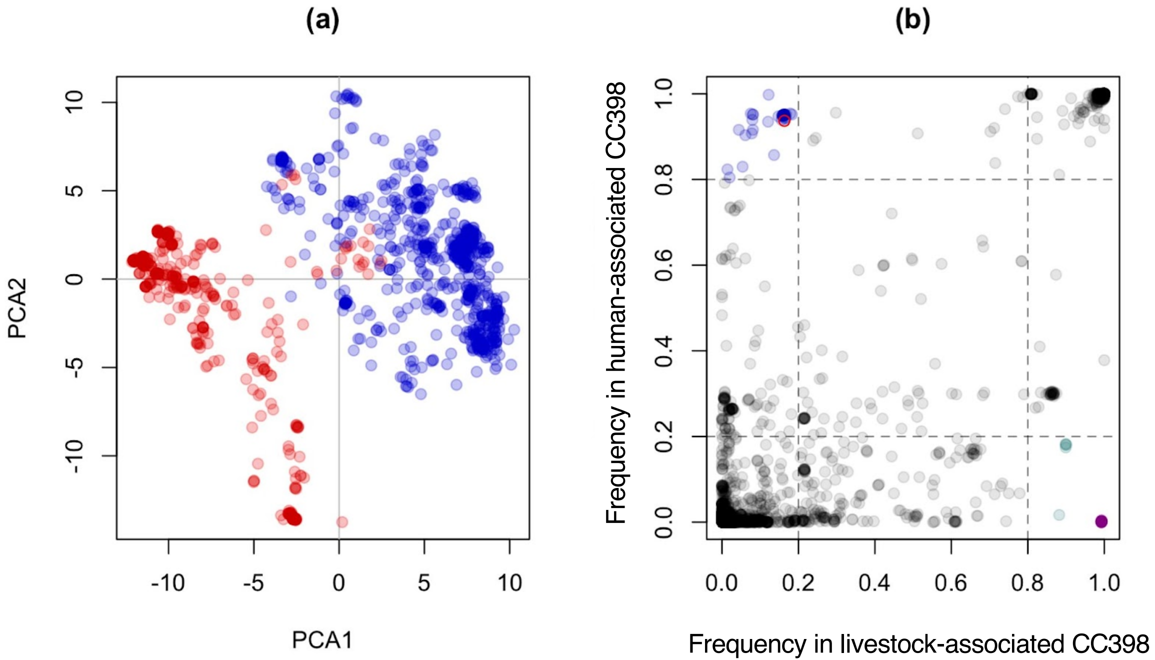

### Figure 2 - Figure Supplement 1

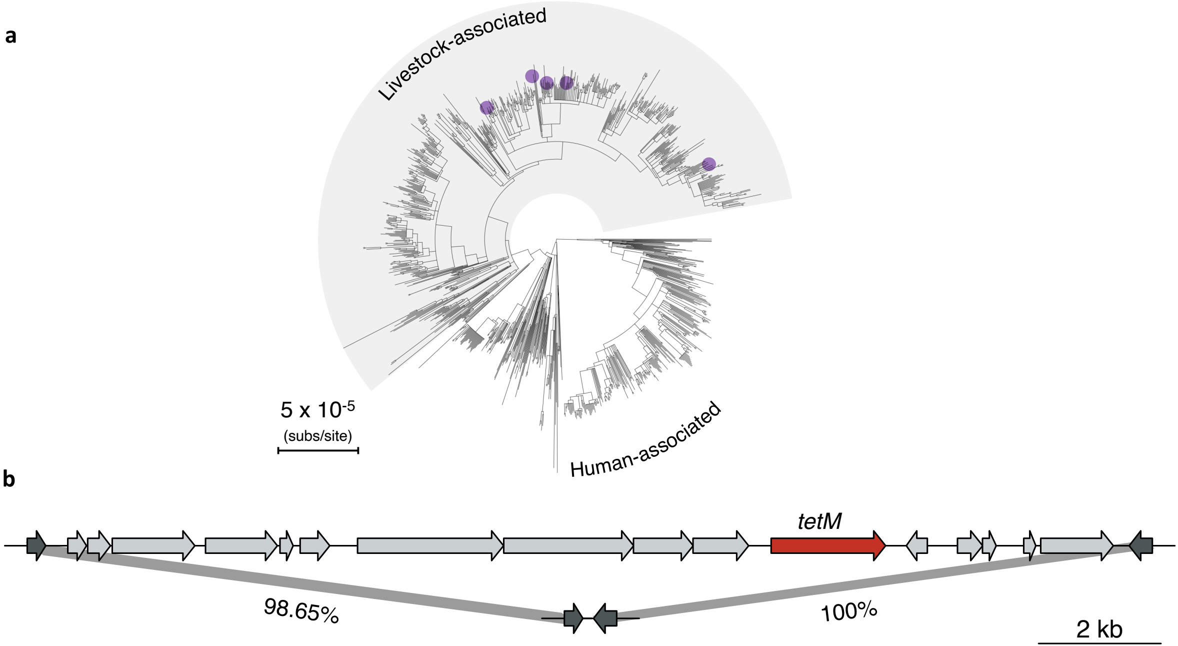

### Figure 3 - Figure Supplement 1

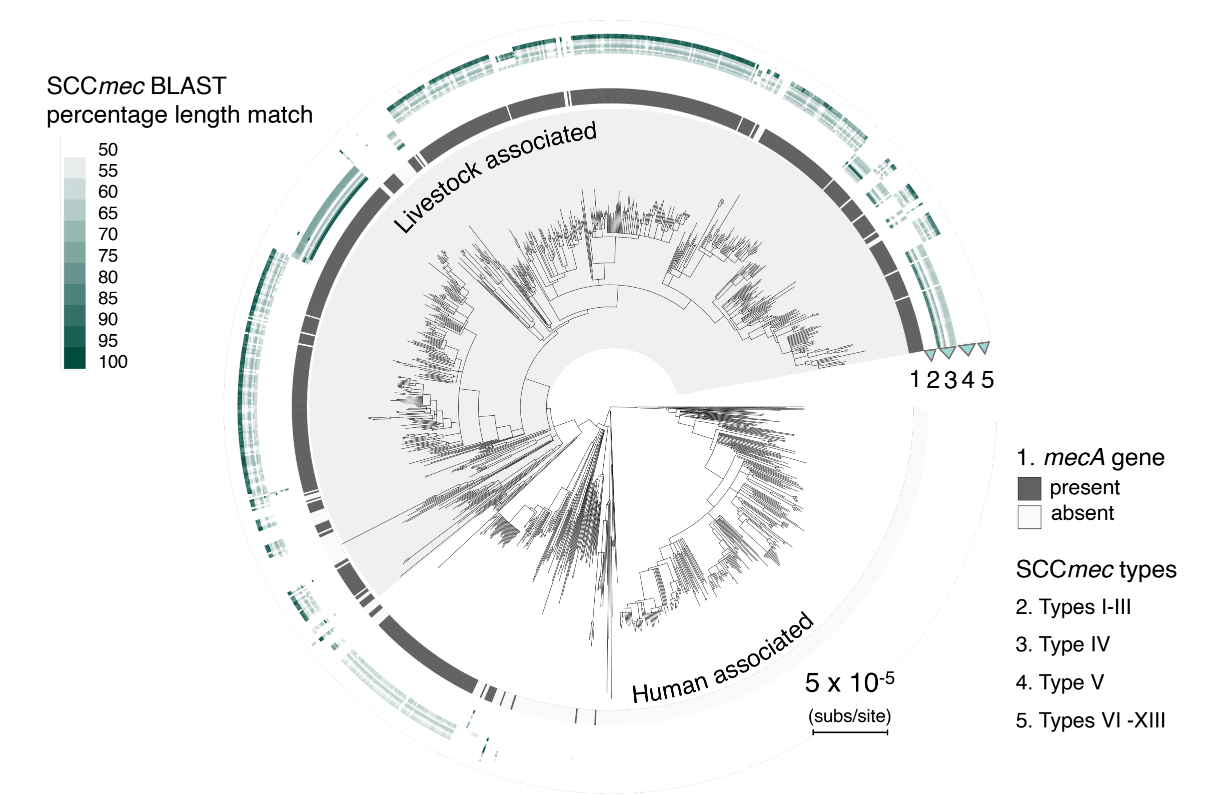

### Figure 3 - Figure Supplement 3

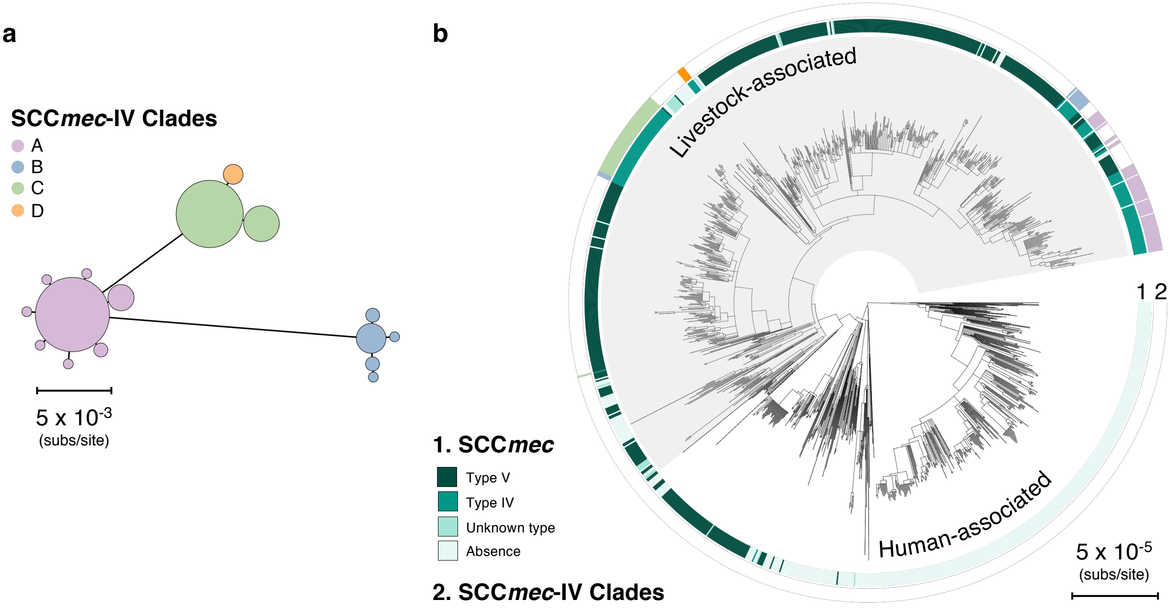

### Figure 3 - Figure Supplement 4

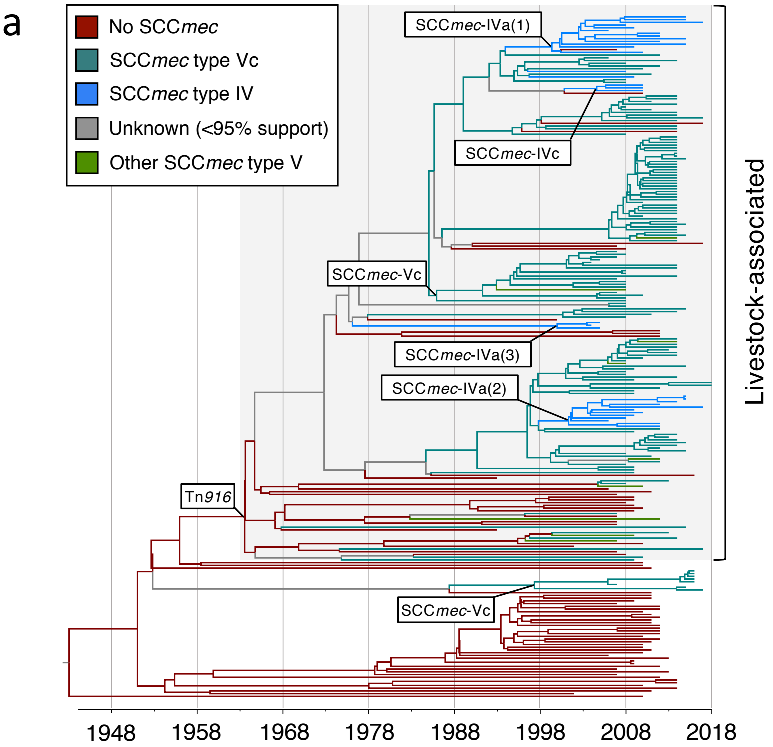

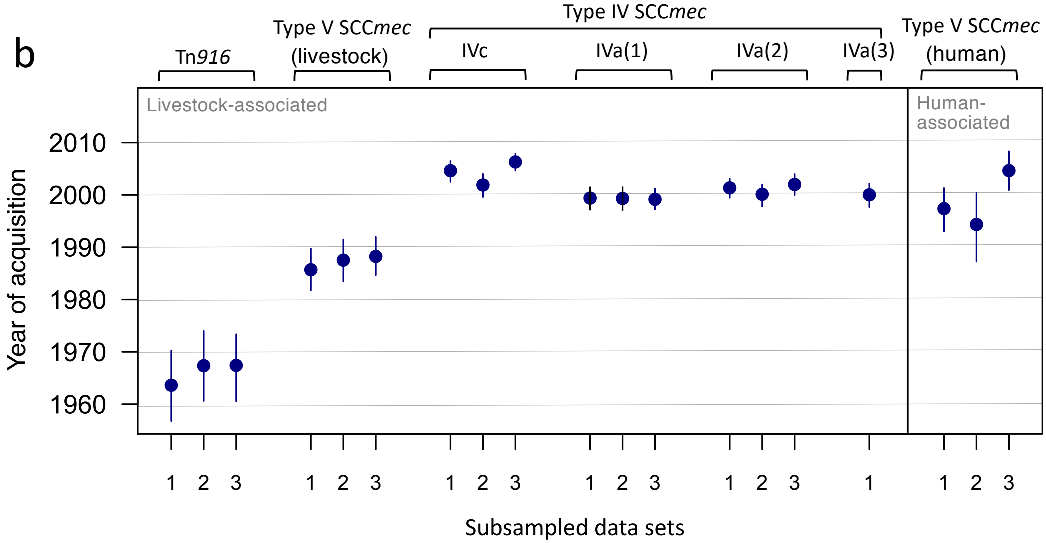

### Figure 4 - Figure Supplement 1

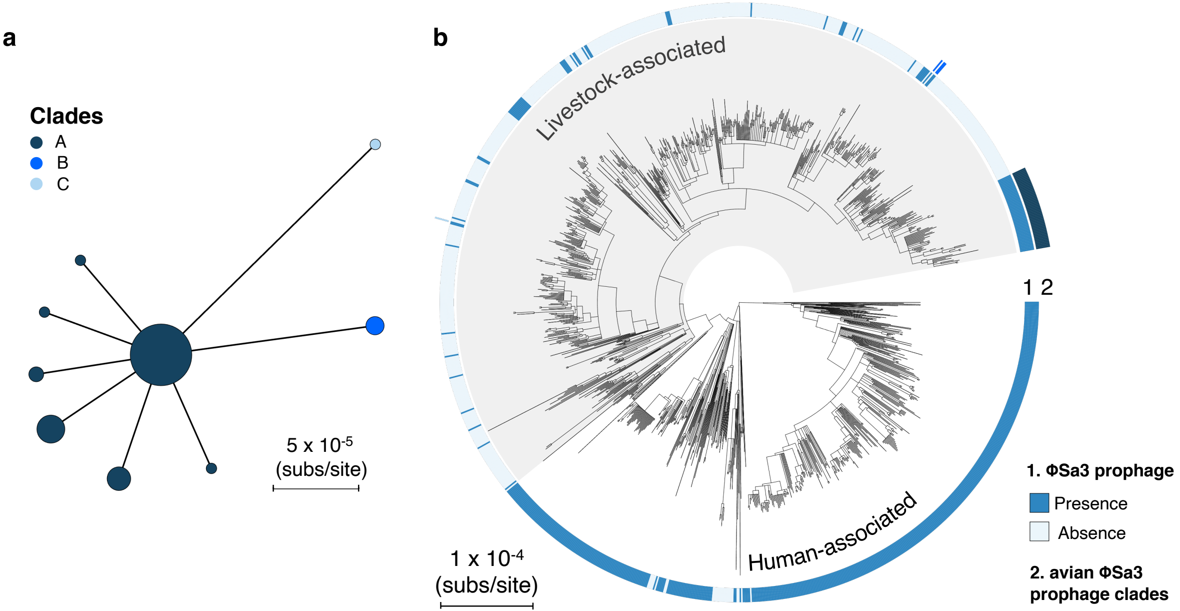

### Figure 4 - Figure Supplement 2

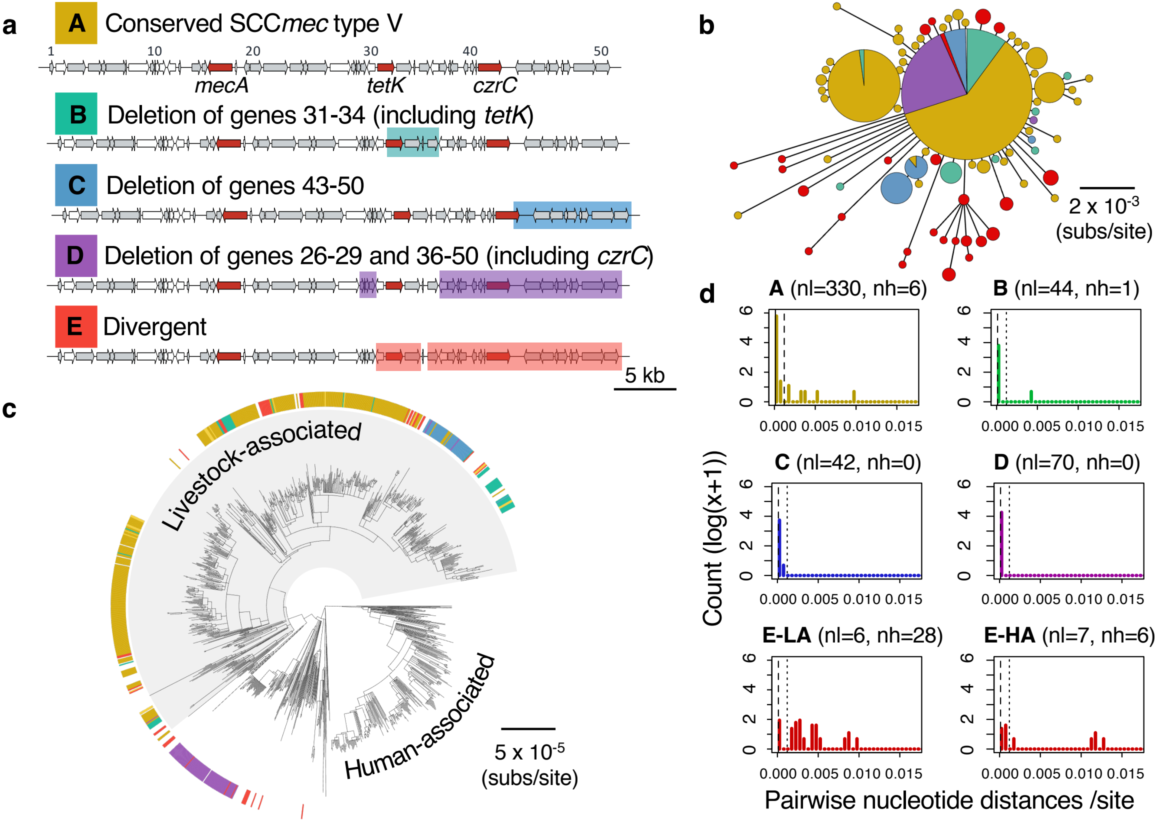

### Figure 5 - Figure Supplement 1

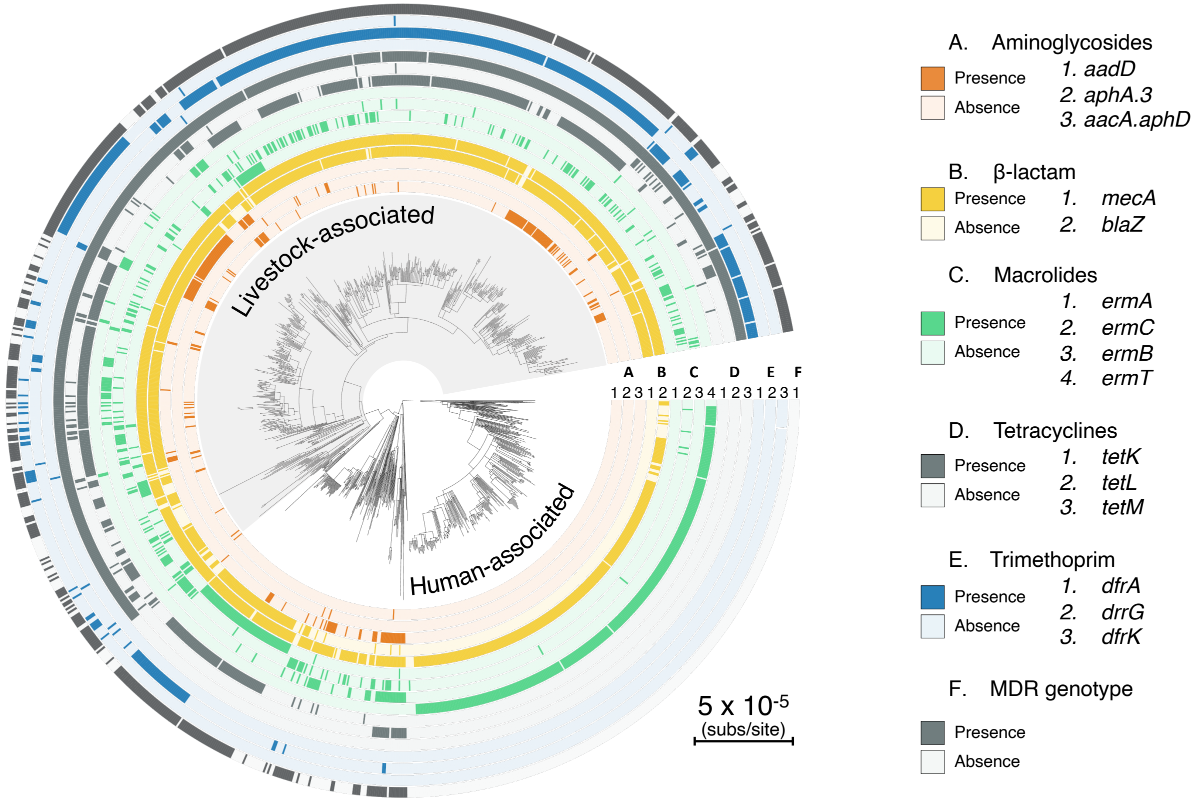

### Figure 5 - Figure Supplement 2

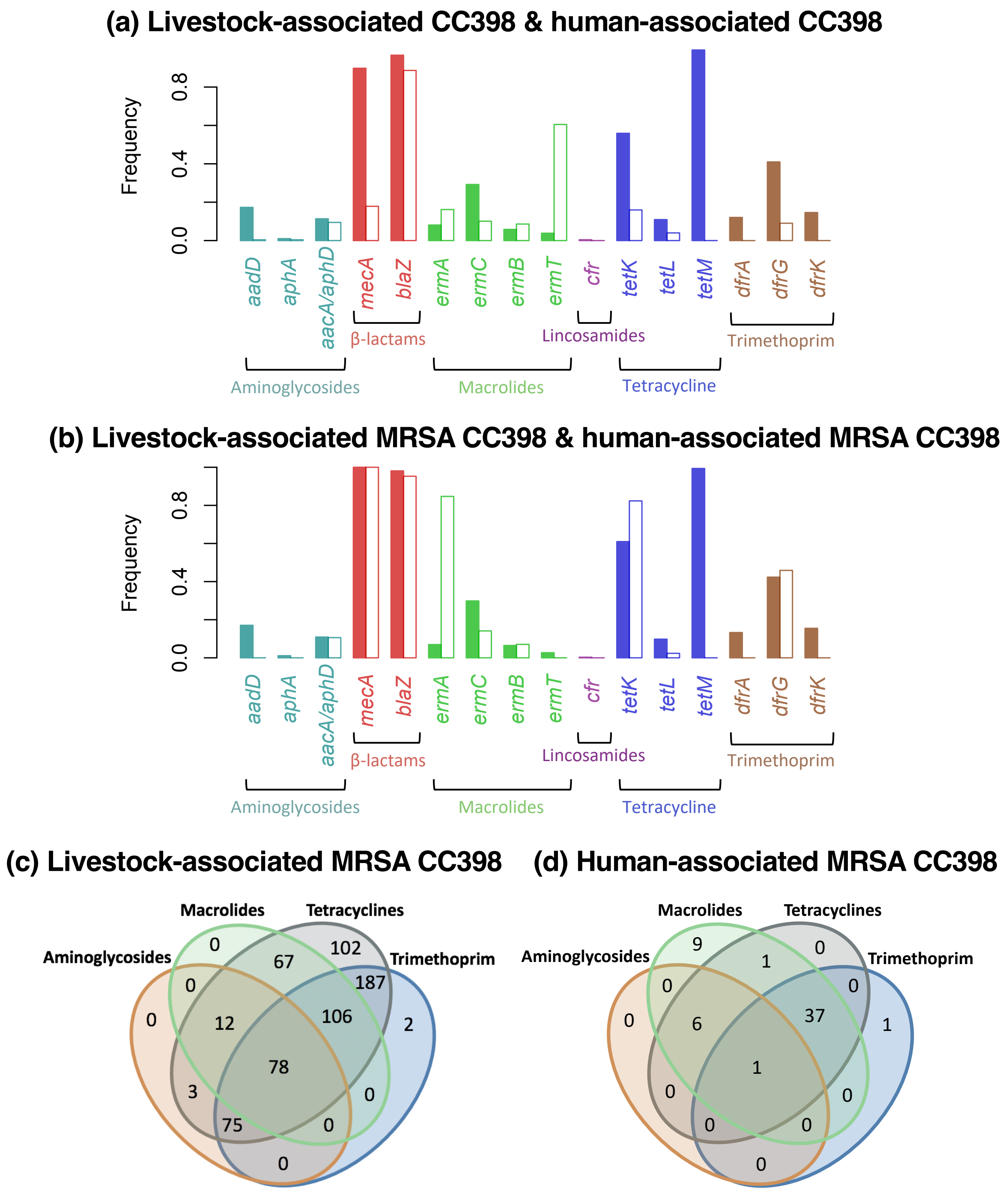

### Figure 5 - Figure Supplement 3

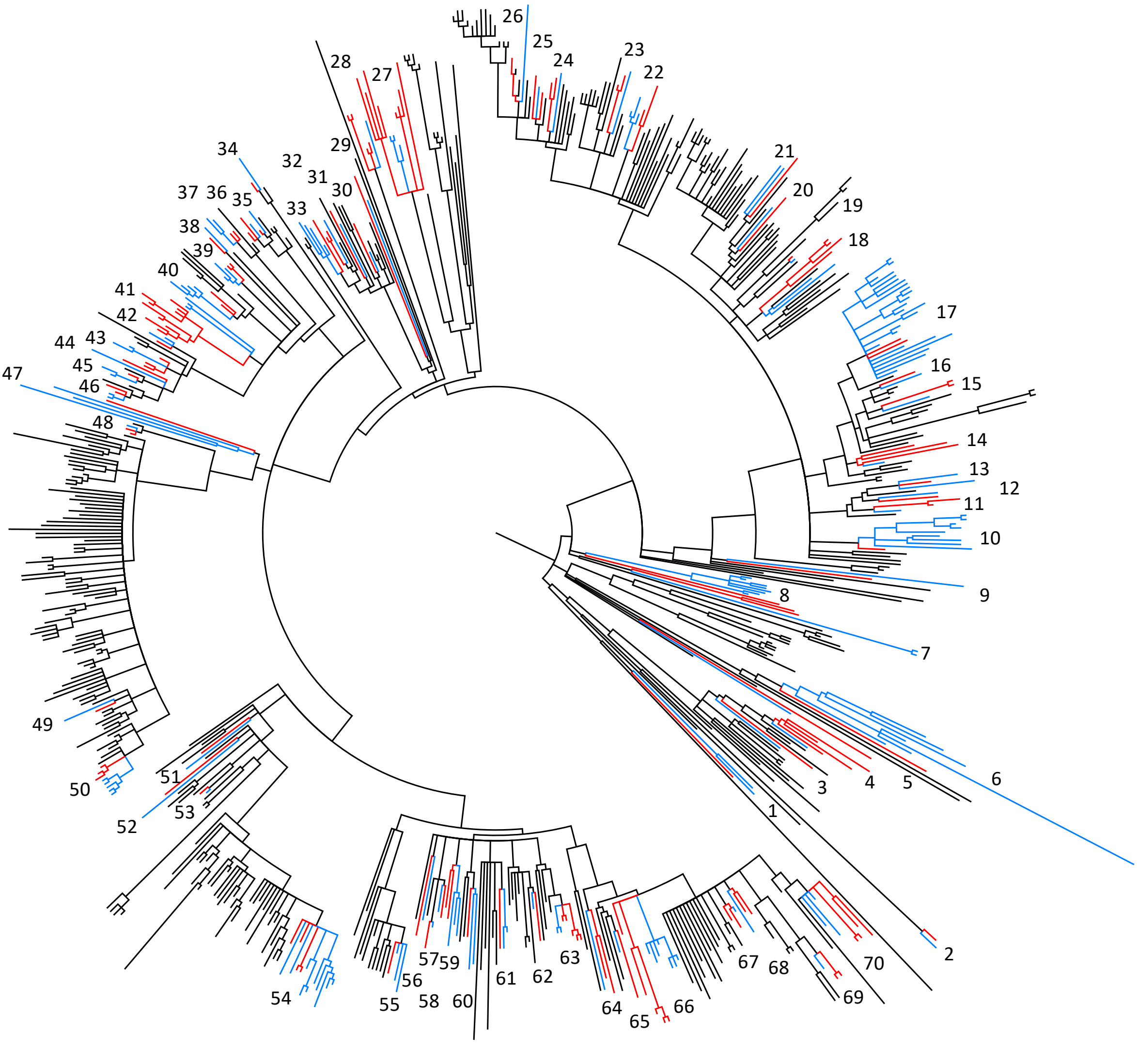
